# Population interconnectivity shapes the distribution and complexity of chimpanzee cumulative culture

**DOI:** 10.1101/2023.08.14.553272

**Authors:** Cassandra Gunasekaram, Federico Battiston, Cecilia Padilla-Iglesias, Maria A. van Noordwijk, Andrea Manica, Jaume Betranpetit, Andrew Whiten, Carel P. van Schaik, Lucio Vinicius, Andrea Bamberg Migliano

**Affiliations:** Department of Evolutionary Anthropology, University of Zurich, Zurich, Switzerland; Department of Network and Data Science, Central European University, Vienna, Austria; Comparative Socioecology Group, Department for the Ecology of Animal Societies, Max Planck Institute of Animal Behavior, Konstanz, Germany; Department of Zoology, University of Cambridge, Cambridge, UK; Institut de Biologia Evolutiva, CSIC - Universitat Pompeu Fabra, Barcelona, Spain; Centre for Social Learning and Cognitive Evolution, School of Psychology and Neuroscience, University of St Andrews, St Andrews, UK; Department of Evolutionary Biology & Environmental Studies, University of Zurich, Zurich, Switzerland

## Abstract

While cumulative culture is a hallmark of hominin evolution, its origins can be traced back to our common ancestor with chimpanzees. Here we investigate the evolutionary origins of chimpanzee cumulative culture, and why it remained incipient. To trace cultural transmission among the four chimpanzee subspecies, we compared between-population networks based on genetic markers of recent migration and shared cultural traits. We show that limited levels of interconnectivity favored the emergence of a few instances of cumulative culture in chimpanzees. As in humans, cultural complexification likely happened in steps, with between-community transmission promoting incremental changes and repurposing of technologies. We propose that divergence in social patterns led to increased between-group mobility in *Homo*, propelling our lineage towards a trajectory of irreversible dependence on cultural exchange and complexification.

**One-Sentence Summary:** Population interconnectivity through migration explains the origins of chimpanzee cumulative culture and why it remained incipient

---

Culture, defined as arrays of behavioral traditions transmitted via social learning (*1*), has been increasingly documented across animal species (*2–5*). Human culture is nevertheless vastly more complex, diverse, and most crucially, cumulative (*6* – *8*). Human cumulative culture provides numerous examples of cultural ratcheting (*8–10*), an evolutionary process that builds cultural traits upon pre-existing ones. Eventually their production may thence become opaque and they can no longer be reinvented from scratch. Cultural ratcheting has only recently been investigated in other species (*11*). Chimpanzees, our closest living relatives, exhibit a vast repertoire of cultural behaviors that vary across sites and are only weakly tied to their habitat (*12–16*). Chimpanzee culture also includes possible instances of cultural accumulation, most evidently in the case of foraging toolsets or multi-step extractive processes (*14*, *17–20*). It remains to be explained what conditions favored the emergence of cultural accumulation in the common ancestor of humans and chimpanzees, and why it remained incipient in chimpanzees in comparison to humans and other tool-making hominins.

Recent analyses have argued that the evolution of a hominin foraging niche, characterized by higher levels of individual mobility and migration played a major role in the cultural differentiation between the two lineages (*21*). Group interconnectivity has been shown to facilitate cultural accumulation in human populations, by preventing erosion of cultural diversity (*22*) and accelerate the recombination of innovations through between-group exchanges (*23*, *24*). Demonstrating the long-term operation of partial connectivity in humans, genetic and archaeological studies have shown that Central African hunter-gatherers have maintained large-scale networks through migration over the past 120,000 years (*25*), as evidenced by varied signs of cultural transmission even between distinct linguistic groups across the whole Congo Basin (*26*).

Investigating levels of interconnectivity between communities may therefore provide relevant insights into cumulative cultural evolution in chimpanzees. Genetic analyses of evolutionary divergence and recent migration among the four chimpanzee subspecies (*27*) revealed a very different pattern of interconnectivity between populations relative to Central African hunter-gatherers. Since the early separation of the four subspecies between 139-633 kya (*28*), low levels of genetic connectivity between the Nigeria-Cameroon, Central, and Eastern subspecies were found during the late Pleistocene and Holocene (*27*). While shared nearly-private rare alleles (NePRAs) provide evidence for reduced connectivity between populations from 15,000 years ago, shared identical-by-descent (IBD) genetic segments suggest some level of migration between Central, Nigeria Cameron and Eastern regions in the past 5,000 years (*27*). Besides being relatively isolated on a regional level, chimpanzees live in large, polygynandrous communities with steep social hierarchies and strict male philopatry, which virtually eliminates male migration (*29*). This contributes to a much lower frequency of between-group exchanges in chimpanzees compared to humans, further limiting cultural diffusion to migrating females (*30*). Hill et al. (*31*) suggest that while an adult hunter-gatherer male may interact with around 300 other tool-making individuals throughout a lifespan, an adult male chimpanzee habitually observes only around 20 others (*31*), or around 40 in the case of females, who typically migrate once in a lifetime.

Given the different social structures of chimpanzees and humans, here we investigate whether the incipient cumulative culture observed in chimpanzees is associated with their relatively modest levels of between-group connectivity and cultural transmission. We combined genetic information on recent between-group migration with chimpanzee cultural data compiled by the Pan African Program (*32*). The latter has built upon earlier surveys (*14*, *16*) to catalogue cultural repertoires in 144 chimpanzee communities across Africa (*12*, *13*), resulting in a growing dataset of community-level cultural traits (*17*, *18*, *33*, *34*). Within this dataset, 35 populations match the NePRA dataset (Fig. 1A, B) and 34 populations match the IBD dataset (Fig. 1C), allowing us to trace group interconnectivity over the past 15,000 years.

**Figure 1.**
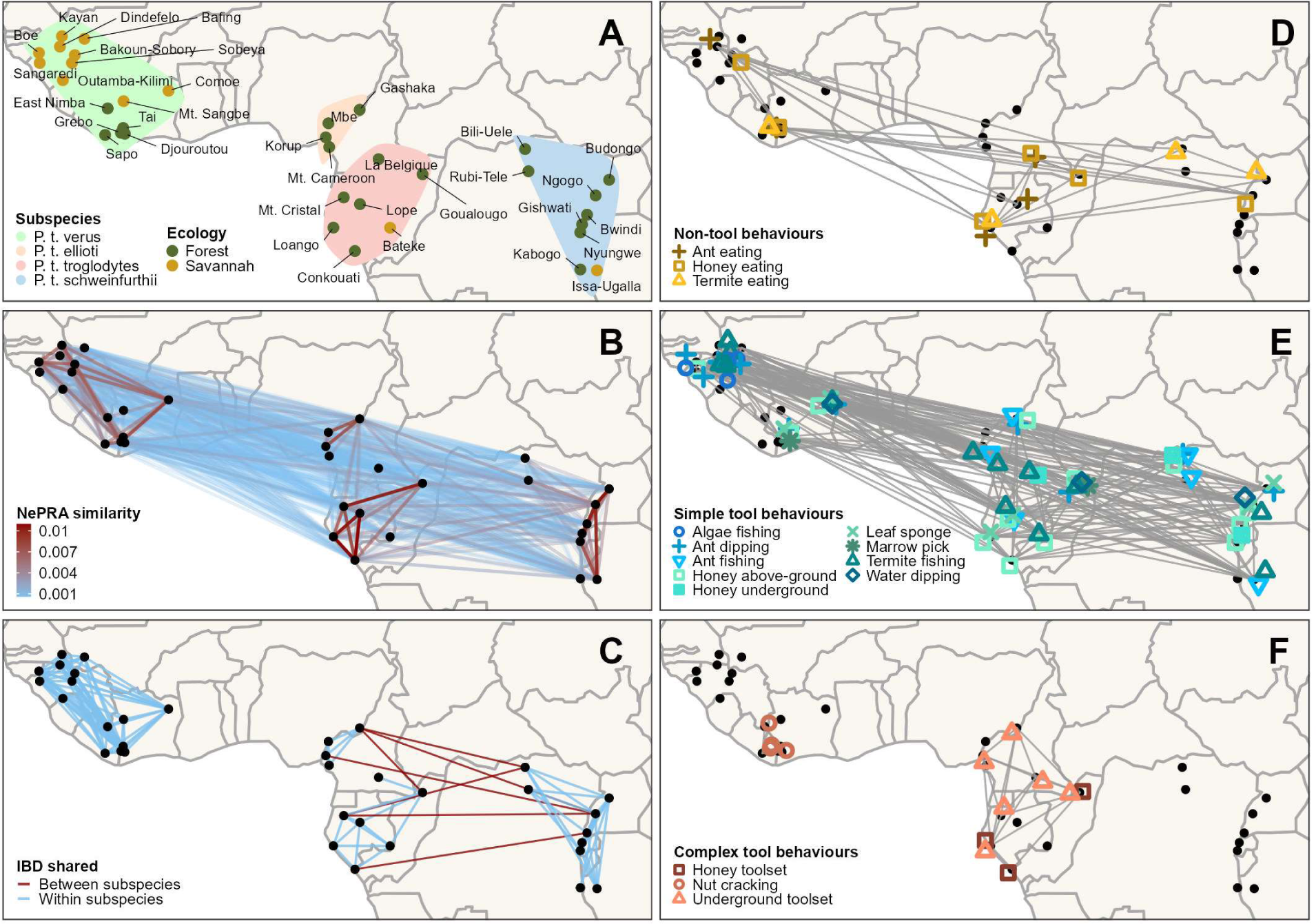
Cultural and genetic connections across chimpanzee populations. Links represent genetic connections based on genetic markers of migration (A-C) or shared knowledge of foraging behaviors (D-F). A) 35 chimpanzee populations included in this study, with habitat and subspecies at each site. B) NePRA connections, with colors indicating proportion of shared NePRAs. C) IBD connections. D) Shared non-tool behaviors. E) Shared simple tool behaviors. F) Shared complex toolset behaviors.

We used the data on recent genetic connectivity to predict the sharing of 15 foraging behaviors between chimpanzee populations. We classified foraging behaviors into three categories of increasing complexity (see SI for details): non-tool behaviors (Fig. 1D), ‘simple’, unitary tool use (Fig. 1E), and ‘complex’ toolset behaviors (Fig. 1F) (*12*, *13*, *35*). We tested the following predictions: (1) foraging without tools and with simple tools should exhibit weak associations with genetic markers of group interconnectivity: they should exhibit a broader distribution across the four subspecies and regions due to a higher likelihood of independent invention at the evolutionary time scale; (2) complex toolsets should be largely explained by genetic markers of recent migration: their distribution should be more dependent on cultural transmission, and therefore more regionally limited due to reduced group interconnectivity in comparison to humans; and (3) as in humans, cumulative cultural evolution in chimpanzees is likely to be a stepwise process facilitated by transmission across multiple individuals, generations and populations.

## Results

### Complex toolsets rely on cultural transmission between populations

To analyze the effect of genetic connectivity on shared knowledge of behavioral traits, we applied Bayesian logistic regression models. We fitted models to predict the dyadic sharing of 15 behaviors (coded as a binary response variable) between populations. We fitted separate models for the sharing of the three behavioral categories (non-tool, simple tool, complex toolset; see SI for details of regressions and full code). As the choice of predictors, all regressions included habitat (savannah or forest) and months of observation at each site as fixed effects; and both type of behavior (e.g. ant eating, termite eating, and honey eating in regressions of non-tool sharing) and population dyadic ID as random intercept effects (see Fig. S3 for random intercept and slope models, which overall render similar results). Each regression included a single genetic or geographic predictor at a time, because genetic markers, region and geographical distances are intercorrelated; see SI for correlation tests).

Genetic markers of recent between-group migration revealed a clear pattern. Dyadic IBD sharing (binary variable; sharing any vs. no IBD segments) had a strong effect on complex toolset sharing (odds ratio (OR) = 2.57, 95% credible interval (CI): [1.15, 5.67]) but no effect on non-tool behavior (OR = 0.67, CI: [0.27, 1.63]), and a weaker effect on simple tool sharing (OR = 1.38, CI: [0.96, 1.96]) (Fig. 2A). NePRA sharing also had the strongest effect on complex toolset sharing (binary: OR = 5.69, CI: [2.14, 17.15], proportions: OR = 3.15, CI: [1.76, 5.72]), a weak effect on simple tool sharing (binary: OR = 1.70, CI: [1.23, 2.33], proportions: OR = 1.44, CI: [1.23, 1.70]), and no effect on the sharing of non-tool behaviors (binary: OR = 1.01, CI: [0.46, 2.23], proportions: OR = 0.93, CI: [0.63, 1.41]) (Fig. 2B, see SI for details). Dyadic region (binary; same vs. different region) supported these results, with same-region populations sharing more complex toolsets (OR = 3.76, CI: [1.75, 8.27], Fig. S2). Geographical distance also confirmed the pattern with a strong negative effect on complex toolset sharing (OR = 0.40, 95% CI: [0.26, 0.59], Fig. S2), suggesting increased sharing of complex toolsets with shorter distances. The associations between genetic markers and complex tool sharing were confirmed by Mantel tests even after controlling for habitat or observation time (Fig. S7). Overall, the results demonstrate a strong association between group interconnectivity and the sharing of complex toolsets.

**Figure 2.**
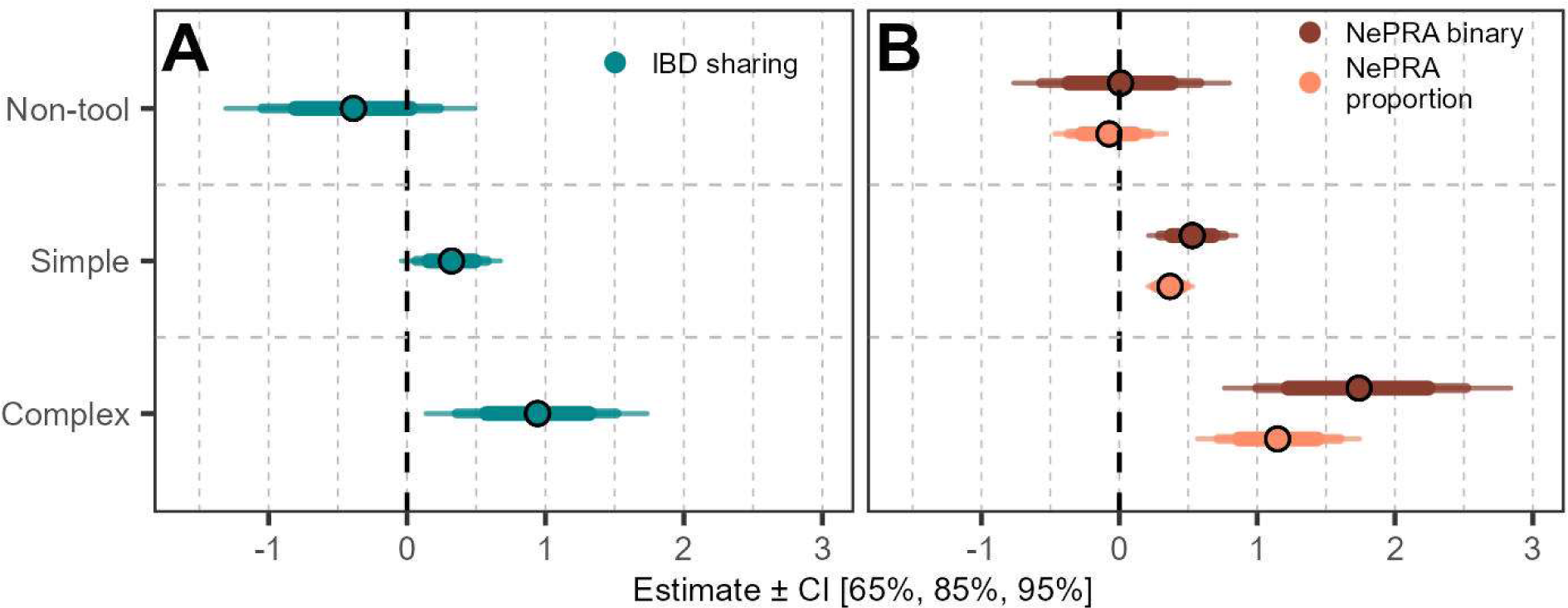
Bayesian logistic regressions of shared cultural traits between chimpanzee populations. Panels display log of the odds ratio estimates and the 65%, 85% and 95% credible intervals. A) Effect of shared IBD segments (binary = sharing any vs. no IBD) on non-tool, simple tool, and complex toolset behaviors. B) Effects of sharing any NePRA (binary; above or below a noise threshold at a similarity of 0.0006, see SI) and proportion of shared NePRAs (see Methods).

### Migration is a key but not sufficient condition for complex toolset sharing

To further quantify the effects of recent migration on non-tool, simple tool and complex toolset sharing, we applied reinforcement analysis (*36*). The technique is derived from social network analysis and compares two networks with the same nodes (in our case, populations) but different structures to determine whether one is likely to be the condition for the existence of the other. Specifically, if the links of network A strongly predict the presence of their corresponding links in network B, then the links of network A are strongly conditioned (or reinforced) by the presence of the links in network B (see SI for further details and examples), providing a possible hint at the directionality of effects between two correlated networks. We built two sets of networks, one with links representing the sharing of a genetic marker (IBD segment or NePRA) between two populations (Fig. 1B, C), and the other with shared cultural behaviors between two populations (non-tool, simple tool and toolset) as links (Fig. 1D-F). We predicted that the sharing of a complex toolset (but not of non-tool or simple tool behaviors) should strongly predict the presence of a genetic marker (as evidence that complex tool sharing requires migration). Conversely, we did not expect a shared genetic marker to strongly predict the sharing of any of the cultural behaviors (since the latter is not a condition for migration, or alternatively, since cultural transmission depends not only on migration but also on other factors, as discussed below).

We first tested whether links representing shared behaviors of increasing cultural complexity predict the presence of an underlying shared IBD segment or NePRA between the same populations. We found that shared complex toolsets predict on average a shared IBD segment with probabilities over 50%, and a strong NePRA similarity in over 80% of cases. Values for non-tool behaviors and simple tools are much lower (Fig. 3A, B), pointing to the dependence of complex toolsets on migration. Conversely, IBD segment sharing, or NePRA sharing (at three increasing levels of similarity) predicted a very small fraction of cultural links at any complexity level (Fig. 3C, D). Together, the results show that at least 80% of the instances of complex toolset sharing are associated with a marker of migration in the past 15,000 years. The values are much lower for simple tools and non-tool behaviors, suggesting that they are more likely to be independently invented. On the other hand, only a small fraction of all genetic links reflecting recent migration between populations results in cultural sharing of any behavioral type. This confirms that the studied behaviors show no signs of genetic determination (*37*), and also indicates that successful cultural transmission through migration may be limited by factors such as the skill levels of migrant females, conformity to community behaviors by low-ranked migrants, or absence of ecological context for expression of a behavior (*30*, *38*).

**Figure 3.**
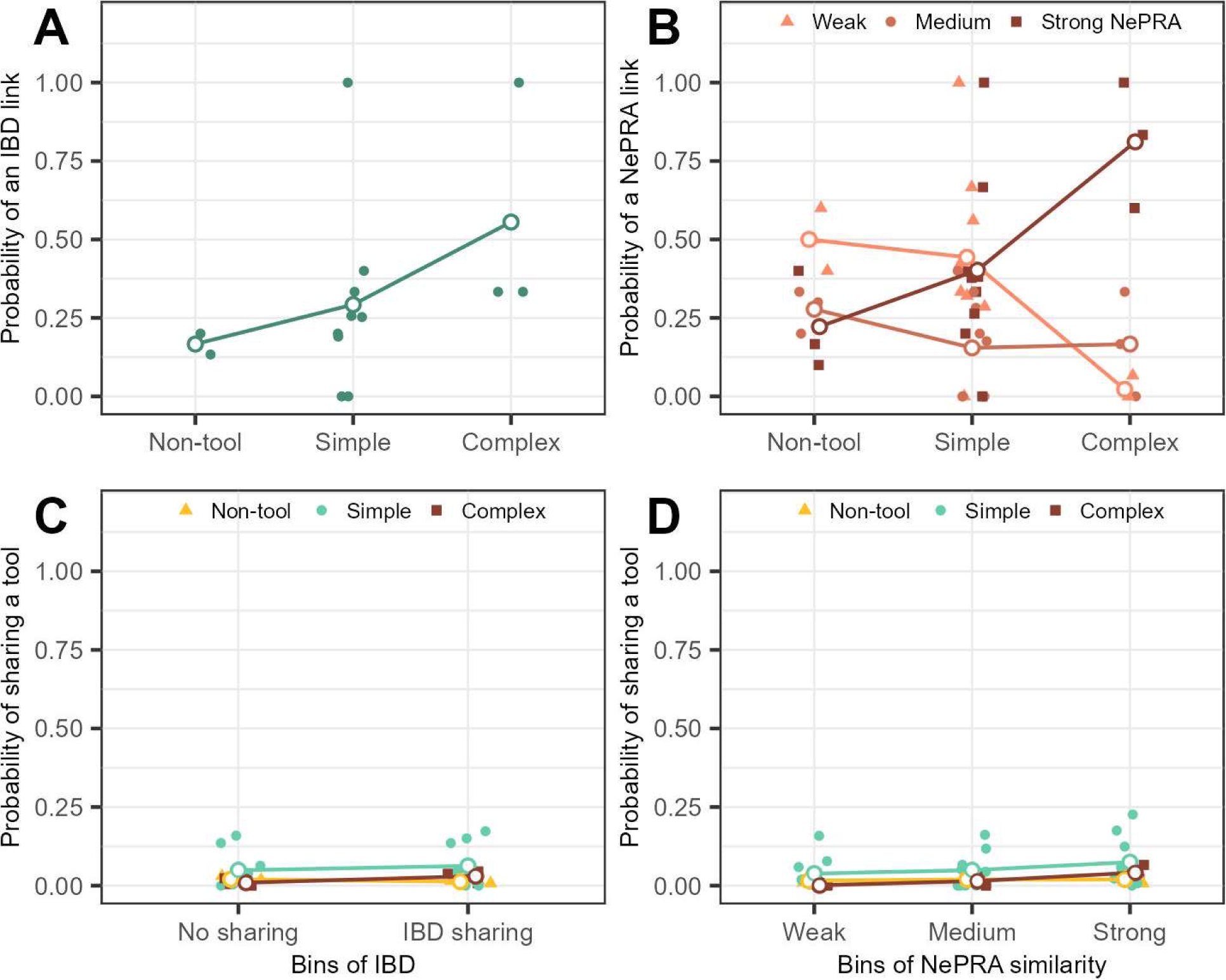
Reinforcement analyses of genetic and cultural sharing networks. A) Probability of the presence of IBD segment links as a function of presence of the corresponding cultural links for the three complexity levels; B) Probability of NePRA sharing at three increasing levels of similarity as a function of cultural links; Panels C) and D) show the probability of presence of links of each behavioral category as a function of IBD links and NePRA similarity links respectively (D). Colored symbols represent the distribution of values of each behavior within each category.

### Evolution of complexity and cumulative culture in chimpanzees

Interconnectivity plays a role in the evolution of cultural complexity in humans by promoting cultural aggregation and recombination (*23*, *24*). We therefore investigated possible signs of stepwise cultural evolution in networks of chimpanzee populations. We assessed evidence of possible evolutionary links between the simple and complex forms of the same underground foraging technology (indicating possible stepwise aggregation), and of repurposing (or major shifts in the use of the technology). Underground foraging tools exhibit distinctive traits of cumulative culture such as variability in number and sequence of tools across populations (*17*, *18*, *34*), and opacity for naive individuals since target resources are hidden from direct observation (*17*). Interestingly, reinforcement analysis produced evidence of a role of migration on complex toolset sharing when all underground toolsets were placed into a single category, suggesting the possibility of an evolutionary link between them (Fig. 3). We compiled all instances of underground foraging in the Central, Nigeria-Cameroon and Eastern regions, including both simple tools (e.g. digging tool) and complex toolsets (e.g. digging stick followed by a dipping tool); exploited resources (honey, termites and ants) from previously published literature (Table S3); and combined this with information on recent genetic links between populations where they are found (shared IBD segments and NePRAs). When we plotted all populations where underground foraging is found and then added their genetic links, we observed that no single population was disconnected from others in the resulting network (Fig. 4A, B). The IBD segment and NePRA networks extend across the three regions through several long links, resembling a pattern often found in hunter-gatherer and other human networks (*26*, *39*).

**Figure 4.**
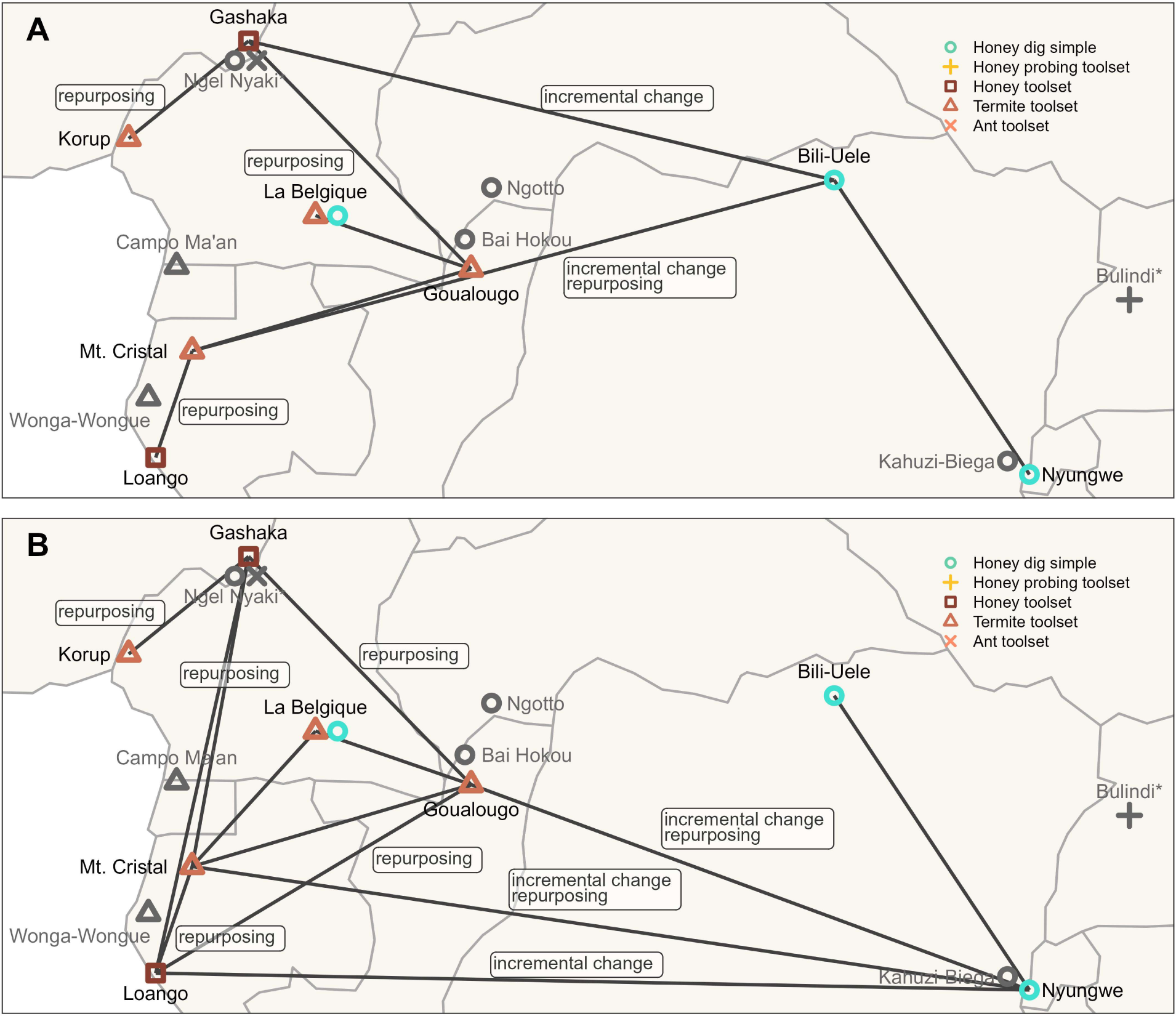
Distribution of underground foraging in chimpanzee populations, underlying genetic links, and putative instances of cultural accumulation. All displayed examples match the description of behaviors in Table S3, apart from the honey probing toolset at Bulindi, which includes a digging stick and an additional investigatory probing stick rather than a honey dipping tool. At Ngel Nyaki, underground dipping tools are reported, but it is unclear whether they are used as single tools or as components of a complex toolset. Lines connect populations sharing (A) IBD segments and (B) strong NePRAs links (similarity > 0.002). Faded, grey populations show cases without genetic data.

This observation further suggests that underground resource foraging may have evolved through incremental changes from use of a single honey digging tool, as found in the East (Bili-Uele and Nyungwe) and in the Central region (La Belgique), into complex toolsets found in the Central and Nigeria-Cameroon regions. For example, Bili-Uele (East), which exhibits only simple underground honey extraction, is directly linked in the shared IBD segment network to Gashaka (Nigeria-Cameroon), where an underground honey toolset is observed, possibly as an example of stepwise aggregation (or addition of a dipping tool to the existing digging tool).

Moreover, in the IBD segment network, the two populations using underground honey toolsets (Gashaka in Nigeria-Cameroon, and Loango in the Central region) are connected to populations displaying complex underground termite foraging such as Korup (Nigeria-Cameroon), Goualougo and Mt. Cristal (Central). This observation and the lack of a simple underground termite foraging variant suggest a possible repurposing of a pre-existing honey toolset for underground termite foraging, implying a major innovation in resource exploitation. Since IBD segments are genetic markers of migration in the past 5,000 years, contact between the three chimpanzee subspecies and tool complexification may have happened recently. Alternatively, the NePRA network tracking a longer evolutionary period (the past 15,000 years) suggests an alternative route of complexification, this time from the simple underground honey tools in Nyungwe (East) to the toolset found in Loango (Central) through incremental addition, followed by repurposing of the technology (from honey to termite foraging) in Goualougo or Mt. Cristal (Central). Although we cannot differentiate between the two possibilities, Mantel tests (Fig. S7D) identified a correlation of underground toolsets with NePRA similarity but not with the more recent IBD segment sharing, pointing to an earlier spread and complexification of the technology within the three subspecies.

In summary, a comparison across sites suggests a variety of markers of cumulative culture in chimpanzee populations: the association between sharing of cultural traits and migration, increased group connectivity and cultural complexity in network positions characterized by increased genetic and cultural exchanges, incremental addition of innovations underlying the origin of complex toolsets, and possibly major innovations with the adaptation of a toolset for new foraging purposes.

## Discussion

In this study we investigated the conditions favoring the emergence of cumulative culture in chimpanzees, and why it has remained incipient in comparison to humans and earlier tool-making hominins. By analyzing cultural and genetic data from 35 populations representing the four chimpanzee subspecies, we showed that group interconnectivity can account to a large extent for differences in the distributions of cultural foraging behaviors. We showed that foraging without tools and with simple tools are weakly associated with genetic markers of recent migration between populations, and are broadly distributed across the four subspecies. We interpret this pattern as a sign that while simple cultural behaviors can be culturally transmitted between populations, they are also likely to be repeatedly invented and reinvented at the evolutionary time scale. We stress that our view is compatible with studies demonstrating the extensive role of social learning in culture (*4*, *40*) and the low probabilities of individual recreation of cultural traits by naive chimpanzees (*9*, *10*, *41*, *42*), since the evolutionary scale of generations and populations provides the opportunity for unlikely events at the individual level to emerge in the long run.

Compared to simple tools, toolsets are less likely to be reinvented due to their complexity and opacity, and accordingly they were strongly associated with genetic markers of migration and between-group transmission in the past 15,000 years. The finding suggests that chimpanzees dispersal patterns in the past may have been sufficient to foster at least an incipient level of cultural accumulation. It is especially interesting that all nut-cracking populations are connected to each other through IBD segment links, pointing to a spread of the technology within the past 5,000 years, matching archaeological evidence confirming the oldest nut-cracking sites at least 4,300 years ago (*43*).

We conclude that cumulative cultural evolution in chimpanzees, as in humans, is a stepwise process facilitated by transmission across multiple individuals, generations and populations. From this perspective, the diversity of complex toolsets in chimpanzees, with nut-cracking limited to Western populations, and honey and termite extraction found in the Central, Eastern and Nigeria-Cameroon regions, results from the gradual and stepwise accumulation of techniques shared across interconnected groups, rather than allopatric differentiation due to isolation. Remarkably, populations that genetically interconnect three chimpanzee subspecies (Mt. Cristal, Gashaka and Goualougo) exhibit some of the highest levels of cultural complexity, further indicating that demographic exchange across regions promotes cultural exchange and complexification. The case study of underground foraging provided possible instances of incremental change as well as major adaptive shifts through repurposing of existing toolsets. We expect that the rich cultural record of chimpanzees may provide many other examples of associations between cultural accumulation and cultural transmission through migration, but unfortunately our analyses are limited to the 35 populations where both cultural and genetic data are currently available. Our study expanded on concepts of between-community cultural transmission proposed in the first systematic analyses of cultural variation in nine chimpanzee communities (*16*) to examine cultural transmission across sites and subspecies, and we hope that in the future a growing dataset of genetic and cultural data will allow for more extensive testing of our predictions.

Finally, the reliance of cumulative culture on between-group interconnectivity also offers a possible explanation for the divergence between chimpanzees and the hominin lineage. The human foraging niche hypothesis (*21*) proposes that cumulative culture has evolved at a faster pace in the ancestral lineage of humans due to the emergence of more extensive social networks in earlier hunting and gathering hominins. The argument points to differences in sociality (*44*), dispersal systems (*45*, *46*), population sizes (*47*), network structures (*24*, *48*), social bonding patterns (*49*), and cooperative breeding (*50*, *51*) which resulted in larger networks and increased connectivity, mobility and cooperation between groups. Archaeological studies indicate that the pattern of population substructuring and interconnectivity may have characterized earlier hominins (*52–54*) and facilitated cumulative cultural evolution. Increased interconnectivity and dependency on cultural exchange may have selected for increased teaching, cooperative learning and joint innovation, setting humans on a unique gene-culture coevolutionary pathway (*55*).

## Acknowledgments

We acknowledge the immense effort of the PanAf consortium to collect and curate the databases. We thank all PanAf-affiliated agents that we list in the SI for generating and making databases freely available in two publications (*12,27*). We thank Susana Carvalho for early discussions during the conceptualization of this project. We extend special thanks to Ammie Kalan, Tomas Marques-Bonet, Martin Kuhlwilm and Claudia Fontsere for their insightful comments and guidance on data analysis. Finally, we thank William McGrew and Thibaud Gruber and for their valued comments on draft manuscripts. We thank Rudy Schlaepfer for the video production.

## Author contributions

Conceptualization: CG, JB, LV, ABM

Data curation; CG

Methodology: CG, FB, CP-I, AM, LV, ABM

Contextualization of analysis: MAvN, AW, CPvS

Formal analysis: CG, FB, LV

Visualization: CG, LV

Funding acquisition: ABM

Project administration: LV, ABM

Supervision: LV, ABM

Writing – original draft: CG, AW, CPvS, LV, ABM

Writing – review & editing: CG, CP-I, MAvN, JB, AW, CPvS, LV, ABM

## Data and materials availability

All data, code, and materials used in the analysis will be made available for purposes of reproducing or extending the analysis on the public repository GitHub upon publication. All data are available will be in the main text or the supplementary materials.

## Supplementary Information

### Materials and Methods

#### Cultural data

To investigate cultural similarities between chimpanzee populations, we used a behavioral database compiled from 144 chimpanzee communities in two previous studies (*12*, *13*). The data for 106 communities was compiled from the published literature and extended with data for 46 communities from the Pan African Program: the Cultured Chimpanzee (for further details see Kalan et al. (*12*), Kühl et al. (*13*)), from which eight were also in the published literature. We extracted communities from the 35 sites that were also in the genetic database and lumped communities at the same site together to match the genetic data. To control for potential ecological factors, we further extracted data on mid-term environmental variability from Kalan et al. (*12*), who classified communities as predominantly savannah woodland or forest (Kalan et al. (*12*), Methods), as well as compiled information on observation time in months at each site (population-average across multiple communities).

We selected foraging behaviors to have a comparative set of behaviors across different complexity levels, and included only behaviors present in at least two populations to compare shared knowledge (Table S1). We stratified behaviors into three groups according to the number of material parts: non-tool, simple (single tool) and complex (multi-part tool). To estimate cultural similarity, we further resolved tool use behaviors according to the type of tools. This was specifically the case for honey foraging, where tools used to extract above-ground or underground substrates differed in the type of tools and in the process of extraction. We divided both the simple honey tool as well as the complex honey toolset into above-ground and underground by screening the published literature from the respective populations. We repeated this procedure for the termite toolset; only one of the 35 populations (Goualougo, Central) had evidence for an above-ground termite toolset (*56*) and was therefore excluded. Due to the similar process of underground extraction using a multipart toolset for honey and termites (*17*), we grouped these into one behavioral category to increase statistical power.

Following Kühl et al. (*13*) and Kalan et al. (*12*), we coded presence of each behavior (1/0) in all 35 populations, so that direct or indirect observations of a behavior at a site were coded as present (1) and otherwise as not observed (0). We excluded cases that were coded as NA (not applicable) by Kalan et al. (*12*). We determined 15 foraging behaviors and classified three as non-tool, nine as simple and three as complex. For statistical analysis of shared knowledge of behaviors, we created a database comprising all possible pairs of the 35 populations. For each behavior we coded shared occurrence (1/0) in all pairs of populations. We coded a behavior as shared (1) if it was present in both populations of a pair. If the behavior was present in only one or none of the populations, we coded not shared (0).

### Genetic connectivity data

We applied two genetic markers to analyze the effect of recent migration on shared knowledge. We used a genetic database from chromosome 21 from Fontsere et al. (*27*), comprising of non-invasive fecal and shed hair samples from 828 individuals at 48 sampling sites across all four chimpanzee subspecies as part of the PanAf Program. Coming from non-invasive samples, only a subset of 240 individuals had high-coverage genotypes (at least 5-fold coverage on average), while the data is patchy and of low coverage for most individuals, as described by Fontsere et al. (*27*).

To assess recent connectivity, we used identical-by-descent-like (IBD) (*26*, *27*, *57*, *58*) segments reflecting a recent common ancestor between two individuals (see details of data processing and analysis in Fontsere et al. (*27*)). For individuals from different sites this implies a recent migration event within the past ∼50 generations. The data included only samples with >5-fold coverage to increase the power to detect IBD-like segments in the limited dataset, resulting in data from 34 sites that match the cultural data (no IBD data from Outamba-Kilimi). We performed all analyses on IBD with these 34 populations. Due to exponential decay of IBD segments through recombination, the length of shared IBD segments is correlated with the time of the migration event, rather than the extent of connectivity between two populations. Therefore, we extracted data on dyadic IBD sharing (binary, sharing any vs. no IBD between population-pairs) from Fontsere et al. (*27*), specifically from the supplementary Table S9. Since the published database did not include IBD-connections between subspecies, we extracted these from Fig. 3 from Fontsere et al. (*27*). The database was further extended by data from a second IBD analysis event by Fontsere et al. (*27*), which included only a subset of the Western populations to control for differences in sampling density (see Fig. S93 and SI in Fontsere et al. (*27*)). Since these two datasets combined were able to identify four additional IBD connections (1. Goualougo (Central) - La Belgique (Central), 2. Comoe (Western) - Mt. Sangbe (Western), 3. Kabogo (Eastern) - Ngogo (Eastern), 4. Nyungwe (Eastern) – Rubi-Télé (Eastern)), we used the combined dataset for further analyses.

To support the analysis of connectivity between populations and increase the time frame of potential genetic and cultural exchange, we further used nearly private rare alleles (NePRAs) (*27*, *59*) which represent potentially earlier migration from one population to another (1.5 kya – 15 kya). We used data from Fontsere et al. based on 963,656 SNPs (on average 26,671 per site) obtained from 434 individuals (on average 11 per site) with at least 1-fold coverage and 38 sampling sites, permissive for missing data (*27*). These SNPs were classified as NePRAs if the allele frequency at a site was larger than zero and the cumulative frequency of this allele was less than one at all other sites. We used a matrix from 35 sites that match the cultural data including binary data (1/0) for each SNP from the GitHub repository from Fontsere et al. (*27*, https://github.com/kuhlwilm/rareCAGA). If a NePRA was detected in at least one sample at a site, the SNP was coded as present (1). If there were no cases of this allele at the site, it was coded as not detected (0). In some cases, a SNP was not sequenced in any of the samples at a given site which was coded as NA. To assess similarity between two populations, we extracted the SNPs that were sequenced in both populations of each pair and calculated the proportion of shared NePRAs. This gives a measure of genetic similarity between populations and potential introgression over the past several thousand years. One limitation of the definition of NePRAs is that populations that have only recently undergone separation, for example in the case of Western chimpanzees who might have emerged from a recent population bottleneck (*27*, *28*), have fewer NePRAs (rare allele variants are at high frequencies in multiple populations) than populations that have diversified earlier and where mutations had more time to occur and spread to other populations. Therefore, genetically close populations might express lower levels of NePRA similarity. Due to probabilistic mutation rate, NePRA similarity representing a very old common ancestor cannot be excluded and is considered as noise. After visual inspection of the distribution of NePRA similarity, we determined a shift in scale at a threshold of NePRA similarity at 0.0006 and classified everything below this threshold as noise (Fig. S1).

### Statistical analysis

To analyze the effect of genetic connectivity on shared knowledge of different behavioral traits, we followed the methods of Kalan et al. (*12*) by applying Bayesian Regression Models (BRMs) with Bernoulli response distribution and logit link function. We conducted all analyses in R (*60*), and implemented the code in RStudio (*61*). We built separate models for non-tool, simple, and complex behaviors, implementing shared occurrence (1/0) of each behavior in the three complexity categories as the response variable.

We fitted an individual model for each of the genetic predictors since NePRA similarity and IBD sharing were highly correlated (Spearman’s rho = 0.645, p-value < 0.001). To analyze whether any exchange of NePRAs or IBD segments could predict shared culture, we fit separate models for binary IBD-exchange (1 = sharing of at least one IBD segment, 0 = sharing no IBD segments) between each population-pair as well as binary NePRA exchange (1/0); NePRA similarity above 0.0006 was classified as potential exchange (1) and below was considered as noise and classified as no exchange (0). To test whether the amount of contact could predict sharing of behaviors, we included the continuous variable of NePRA similarity. Due to a high correlation with genetic predictors, subspecies (Spearman’s rho between NePRA similarity and subspecies = 0.660, p-value < 0.001, Spearman’s rho between IBD and subspecies = 0.793, p-value < 0.001) and geographic distance (Spearman’s rho between NePRA similarity and distance = −0.597, p-value < 0.001, Spearman’s rho between IBD and distance = −0.623, p-value < 0.001), we performed two separate models with either same subspecies (binary, 1 = same subspecies, 0 = different subspecies) or geographic distance in km calculated from coordinates at the center of the sites (*12*, *13*, *27*) between a pair of populations (Fig. S2). To control for variation in observation time at different sites, we extracted the shorter observation time in months among the population-pair and included it as a fixed effect (shared observation time). Additionally, we included the binary variable of same habitat (1 if both populations in a pair have the same habitat / 0 if the habitat from the two populations are different). We added the population-pair and behavior as random effects (*62*). Due to the small sample size, we did not include random slope effects. However, we did estimate random slopes of behavior on the genetic or geographic predictor, observation time and habitat (Fig. S3). After inspection of the distribution of the continuous predictors, we log-transformed geographic distance and NePRA similarity and z-transformed both (β ∼ *N*(0,1)) (*62*). Following Kalan et al., we also z-transformed shared observation time. Since additionally log-transforming the shared observation time yielded similar results, we continued analysis with only z-transforming shared observation time (Fig. S4).

We used the function *brm* from the R-package *brms* (version 2.18.0) (*63*) to fit all BRMs. By default, we ran 2,000 iterations including 1,000 iterations for warm-up over four MCMC chains, creating 8,000 usable posterior samples. Model diagnosis of each model showed that there were no divergent transitions after warm-up; all MCMC chains were stationary and converged and all *Rhat* values were below 1.01 (*64*). We used weak priors with a standard normal distribution with a mean of 0 and a standard deviation of 1. To assess the effect of the priors, we also tested a wide prior (β ∼ *N*(0,10)) (Fig. S5). We reported the odds ratios of the mean estimate to compare the effect sizes of the predictors on each of the samples, and assessed the significance of the results using the credible intervals at 65%, 85% and 95% (mean estimate of the marginal posterior distribution, credible intervals around the mean estimate of the posterior distribution) (Fig. 2, table S2). We tested the association between IBD and shared culture using a combined dataset from two IBD sequencing events (the first including all populations and the second using a subset of the first data); however, we checked how the first dataset performed individually by coding IBD-connections identified only by the second sequencing event (Fig. S93, Fontsere et al. (*27*)) as 0 (Fig. S6).

We applied Mantel tests to assess correlations between cultural, geographical, ecological and genetic distances. We calculated Jaccard distances between population-pairs for non-tool, simple and complex using the *vegdist* function from the *vegan* R-package (version 2.6.2) (*65*) and estimated behavioral similarity between population-pairs for each of the three samples by subtracting the Jaccard distance from 1 for more intuitive analysis of the results. We further applied the NePRA similarity and IBD sharing matrices as measures of genetic similarity, and geographic proximity by dividing 1 by distance in km. Additionally, we used a matrix of same habitat (1/0) as well as same subspecies (1/0) and shared observation time (continuously increasing). We performed Mantel tests using the *ecodist* R-package (version 2.0.9) (*66*) with the Spearman correlation coefficient between each pair of similarity/proximity matrices being calculated from 500 resamples. We performed simple Mantel tests in which each pair of matrices was tested (Fig. S7A), as well as partial Mantel tests controlling for same habitat (Fig. S7B) as well as shared observation time (Fig. S7C). Further, we performed a separate simple Mantel test correlating each complex behavior with either IBD sharing or NePRA similarity (Fig. S7D). When interpreting these results, the low sample size within each complex behavioral category and the following weak statistical power of these results must be considered and compared with other analyses.

### Social network analysis

To investigate the relationship between the cultural and genetic transfers resulting from between-group migration, we applied reinforcement analysis (*36*). The technique attempts to determine the directionality of effects between two networks with similar nodes (in our case, populations) but different links, by reciprocally predicting the presence of a link in a network from the strength of the same link in the other. As an example, let us imagine a ‘marriage’ network M where a link represents a married couple, and a ‘lifespan’ network L with the same nodes, but where a link is present if the two individuals were alive at the same time. In this case, links in the marriage network will always predict the presence of the same link in the lifespan network, since being married is conditioned upon the two individuals being contemporaneous. Reversely, links in the lifespan network will predict marriage links with much lower probability, since being contemporaneous is not conditioned upon being married, or alternatively, since many factors other than being alive at the same time are needed for marriage to happen. The rationale applies to networks characterized by links categorized at distinct levels. For example, assume a network C including only married couples with two link levels; no children, or with children. A second network T includes links representing time of marriage; less than six months, or over six months. We expect a dramatic increase in the proportion of ‘over six months’ links predicted by ‘with children’ links (since pregnancy length is nine months). Conversely, the increase in the proportion of predicted ‘with children’ links by ‘over six months’ links is expected to be less pronounced, since there are many childless couples married for longer than six months (due to factors other than duration of marriage behind the decision to have children).

We created an individual network of shared knowledge for each behavioral type, where a link represented sharing the respective behavior. Next, we divided the IBD network into sharing-IBD and no-sharing, and the NePRA network into three levels of genetic similarity. The first bin (weak links, N = 322) contained all links that had a NePRA similarity < 0.0006 (below our defined threshold for noise). The second and third bin were divided at 0.002, creating two almost equally sized bins, with the second bin containing links of NePRA similarity between 0.0006 and 0.002 (medium links, N = 137), and the third bin comprising all links with a similarity larger than 0.002 (strong links, N = 136). The first bin in the analysis involving IBD length contained population-pairs that did not share any IBDs (no sharing, N = 428), while the second bin contained links that shared at least one IBD (IBD sharing, N = 133). To analyze the directionality of the association, we calculated the probability of links in each genetic subnetwork that also displayed a link in each of the behavioral networks. To test for reverse directionality, we calculated the proportion of links in each bin of the two genetic networks for each of the behavioral network, and reported averaged results across behaviors for non-tool, simple and complex (Fig. 3A-D).

### Visual inspection of distribution of cultural traits

To examine how various forms of a technology may relate to each other and with patterns of interconnectivity, we summarized all simple and complex instances of underground foraging in the Central, Nigeria-Cameroon and Eastern regions using all 107 populations from the compiled database from Kühl et al. and Kalan et al. (Fig. S8) (*12*, *13*). We divided the behaviors according to the resources extracted (honey, termites and ants) as well as the different steps by extending the database by screening previously published data (Fig. 4, Table S3). We classified the increase in complexity of a variant of the behavior at a site into incremental change (adding an additional tool) and repurposing (using the toolset for a new substrate) (Table S3). We analyzed the genetic links (IBD: Fig. 4A, NePRA: Fig. 4B) between the respective populations for which we had genetic data and identified the changes in complexity between these populations.

## Supplementary Text

### Extended list of acknowledgements

We thank the following members of PanAf the cultured chimpanzees, for making their datasets publicly available in two different publications: A. K. Kalan, L. Kulik, M. Arandjelovic, C. Boesch, F. Haas, P. Dieguez, C. D. Barratt, E. E. Abwe, A. Agbor, S. Angedakin, F. Aubert, E. A. Ayimisin, E. Bailey, M. Bessone, G. Brazzola, V. E. Buh, R. Chancellor, H. Cohen, C. Coupland, B. Curran, E. Danquah, T. Deschner, D. Dowd, M. Eno-Nku, J. M. Fay, A. Goedmakers, A.-C. Granjon, J. Head, D. Hedwig, V. Hermans, K. J. Jeffery, S. Jones, J. Junker, P. Kadam, M. Kambi, I. Kienast, D. Kujirakwinja, K. E. Langergraber, J. Lapuente, B. Larson, K. C. Lee, V. Leinert, M. Llana, S. Marrocoli, A. C. Meier, B. Morgan, D. Morgan, E. Neil, S. Nicholl, E. Normand, L. J. Ormsby, L. Pacheco, A. Piel, J. Preece, M. M. Robbins, A. Rundus, C. Sanz, V. Sommer, F. Stewart, N. Tagg, C. Tennie, V. Vergnes, A. Welsh, E. G. Wessling, J. Willie, R. M. Wittig, Y. G. Yuh, K. Zuberbühler, H. S. Kühl, C. Fontsere, M. Kuhlwilm, C. Morcillo-Suarez, M. Alvarez-Estape, J. D. Lester, P. Gratton, J. M. Schmidt, T. Aebischer, P. Álvarez-Varona, A. K. Assumang, D. Barubiyo, A. Carretero-Alonso, A. Dunn, J. Dupain, V. E. Egbe, O. Feliu, R. A. Hernandez-Aguilar, I. Imong, M. Kaiser, M. Kambere, M. V. Kambale, A. Laudisoit, M. Llorente, F. Mulindahabi, M. Murai, S. Nixon, C. Orbell, L. Riera, L. Sciaky, L. R. Tédonzong, E. Ton, J. van Schijndel, V. Vergnes, R., K. Yurkiw, J. Hecht, L. Vigilant, A. M. Andrés, D. A. Hughes, E. Lizano, T. Marques-Bonet.

**Fig. S1.**
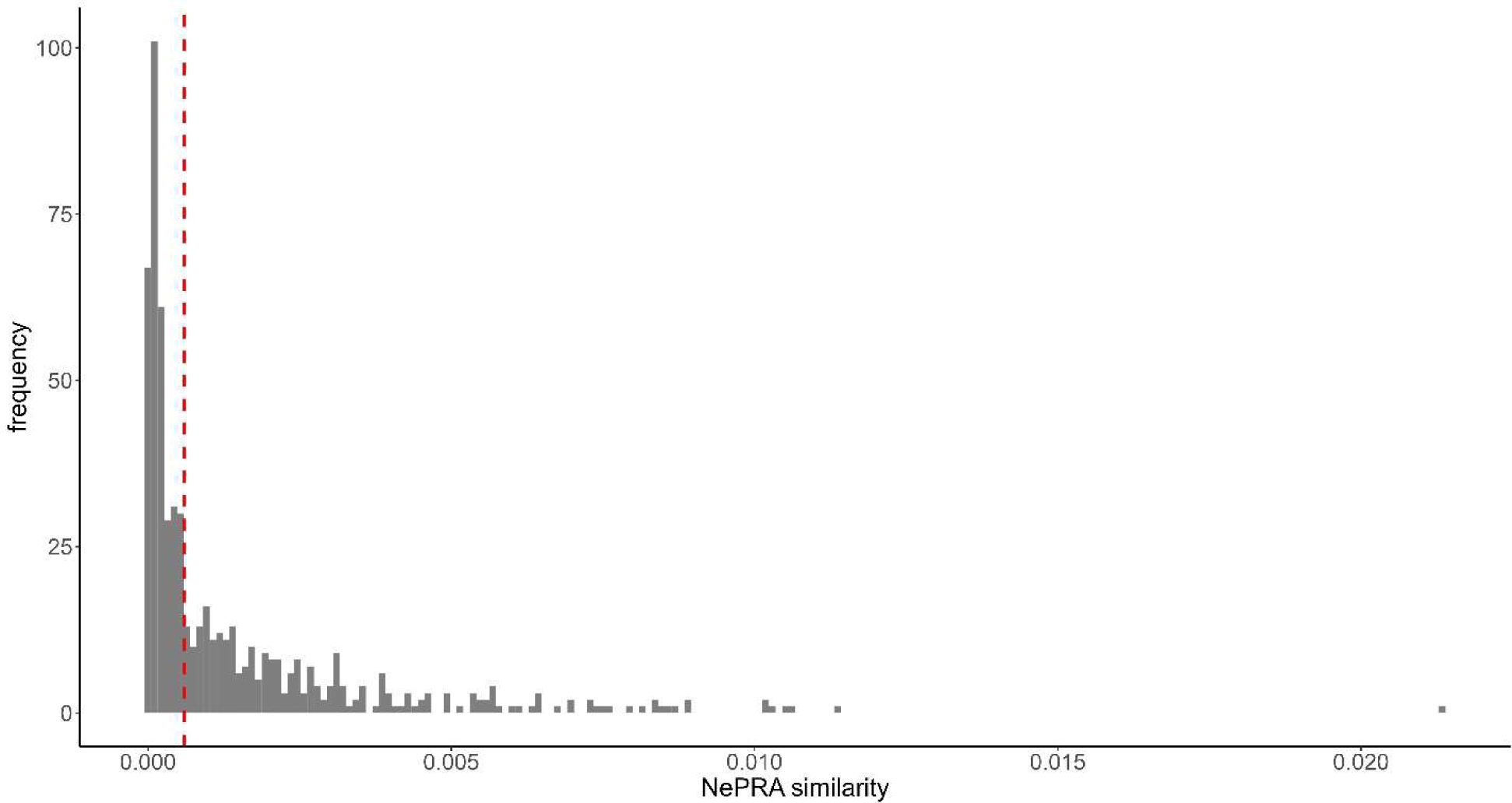
Distribution of NePRA similarity. The red dashed line indicates the noise-threshold at 0.0006. The maximum NePRA similarity is at 0.021 between Conkouati (Central) and Loango (Central).

**Fig. S2.**
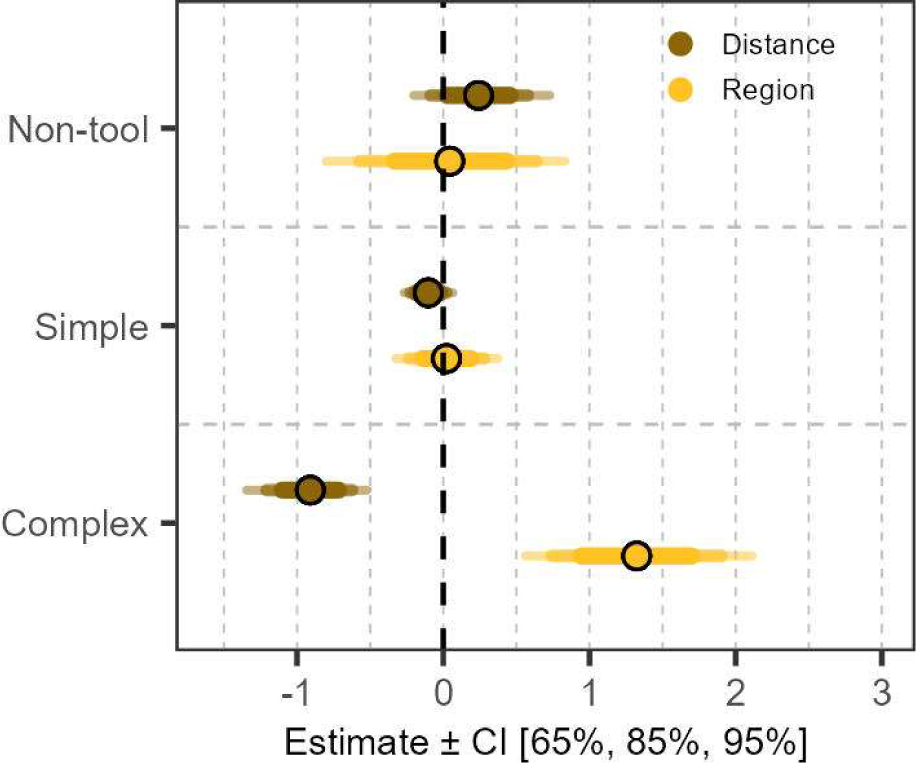
Bayesian logistic regression of effects of geographic distance and same region (1/0) on shared cultural traits between chimpanzee populations. The plot shows the mean of the marginal posterior distribution (dots) and the 67%, 87%, 97% credible intervals centered on the mean (line thickness corresponds decreasingly with size of credible intervals).

**Fig. S3.**
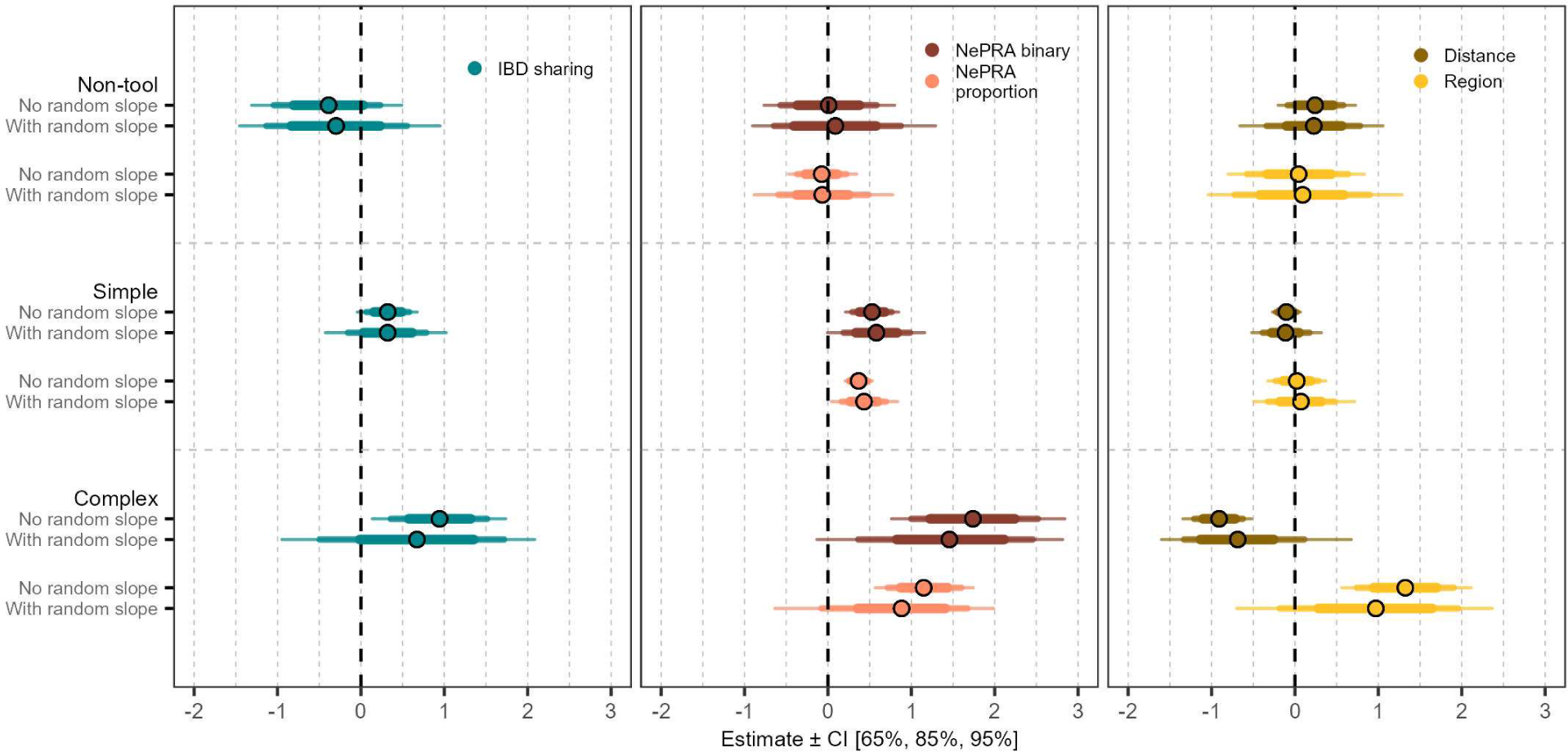
Analysis of random slope effects of behavior. As expected, credible intervals are wider when including random slope effects, but the mean estimate does not differ largely. The plot shows the mean of the marginal posterior distribution (dots) and the 67%, 87%, 97% credible intervals centered on the mean (line thickness corresponds decreasingly with size of credible intervals).

**Fig. S4.**
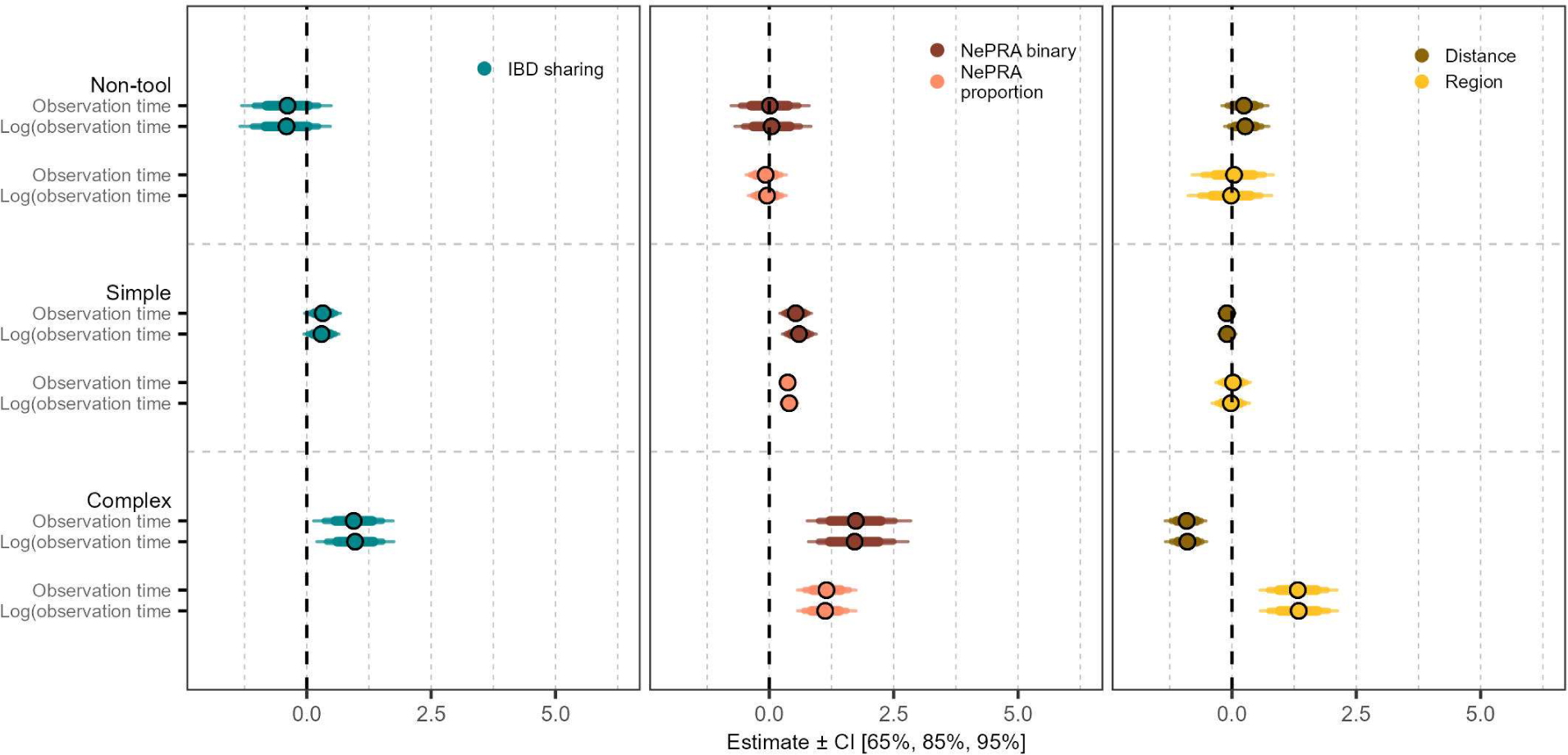
Analysis of log transformation of observation time. The graph shows that the log transformation of shared observation time has no clear effect on the outcome. The plot shows the mean of the marginal posterior distribution (dots) and the 67%, 87%, 97% credible intervals centered on the mean (line thickness corresponds decreasingly with size of credible intervals).

**Fig. S5.**
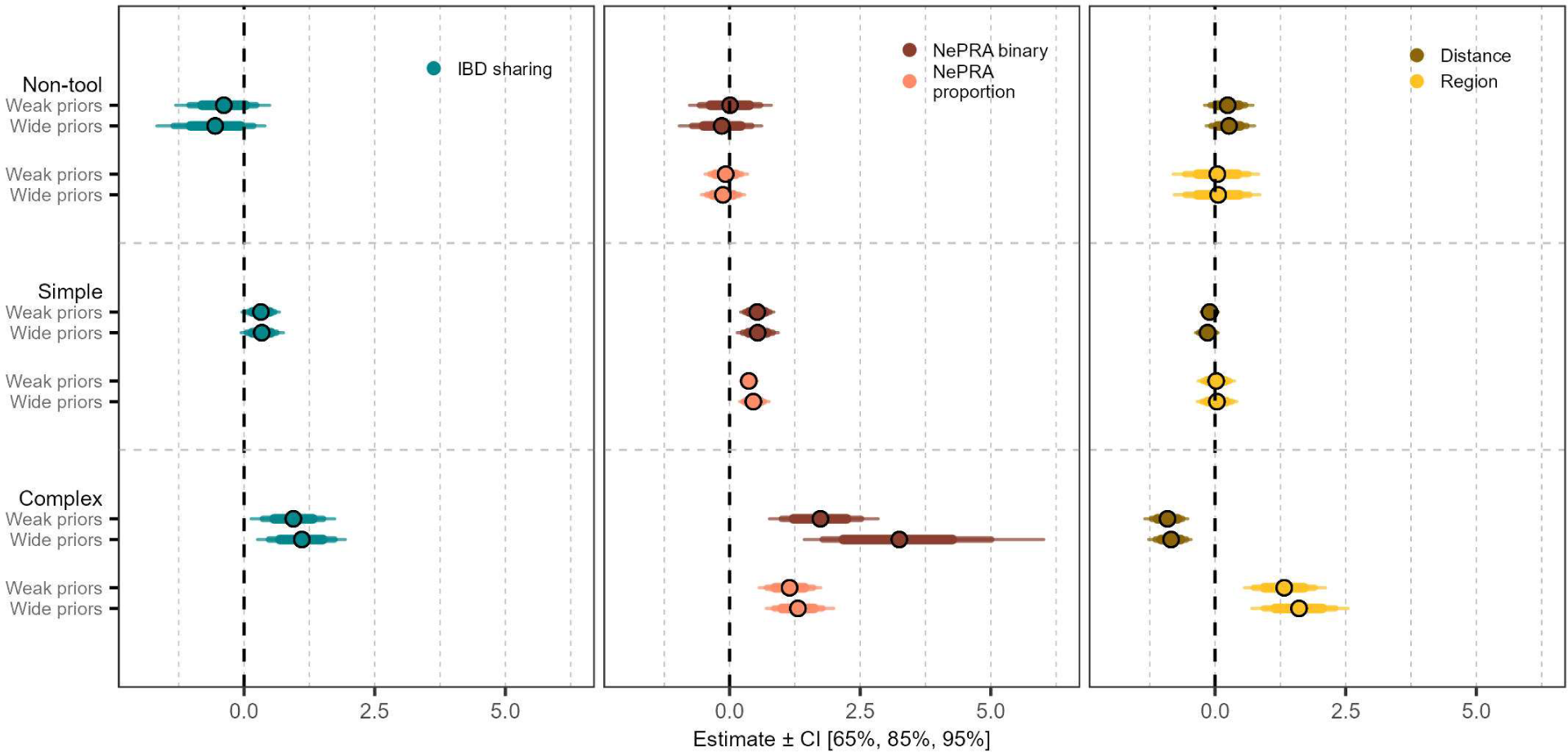
Analysis of prior sensitivity. Interestingly, only the models testing complex tools seem to vary when implementing a wide prior (*N*(0,10)) compared to a weak prior (*N*(0,1)), specifically for the NePRA binary predictor where the distribution is wider. Still the mean estimates differ minimally. The plot shows the mean of the marginal posterior distribution (dots) and the 67%, 87%, 97% credible intervals centered on the mean (line thickness corresponds decreasingly with size of credible intervals).

**Fig. S6.**
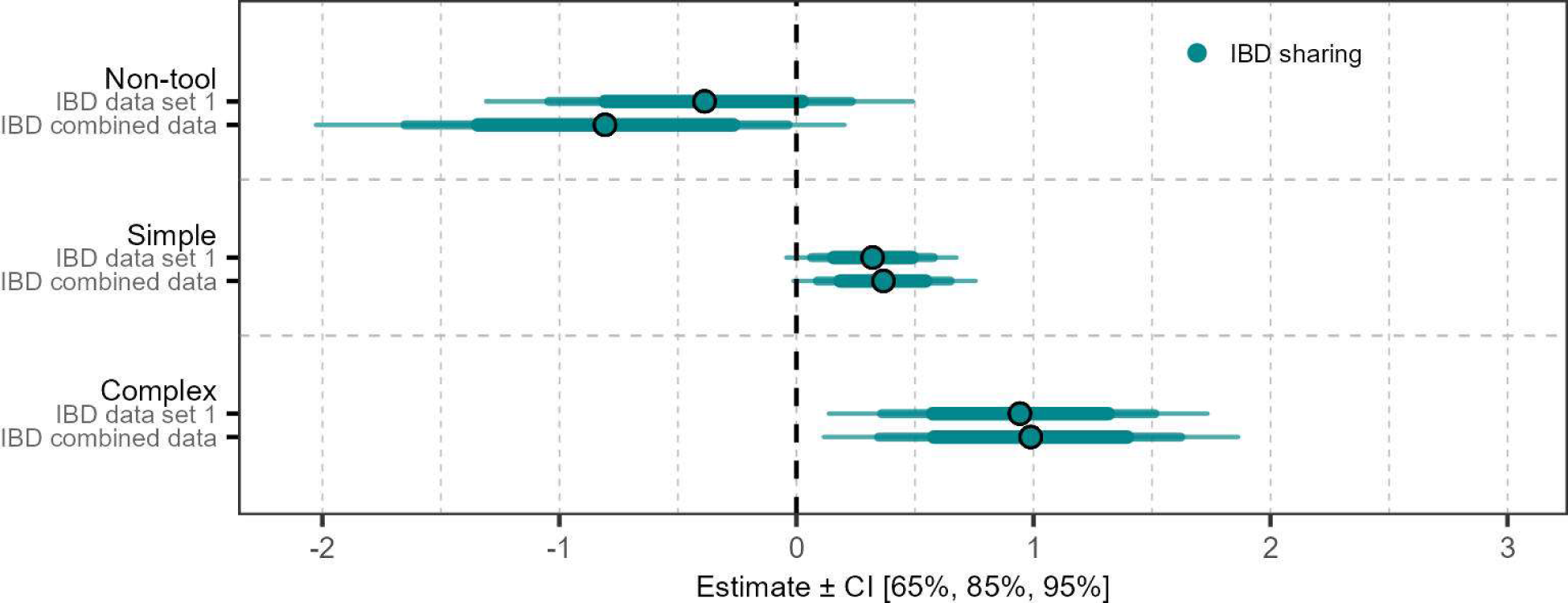
Analysis of robustness of IBD data. As expected, there is no clear difference in results between the IBD data set 1 and the combined data set. The plot shows the mean of the marginal posterior distribution (dots) and the 67%, 87%, 97% credible intervals centered on the mean (line thickness corresponds decreasingly with size of credible intervals).

**Fig. S7.**
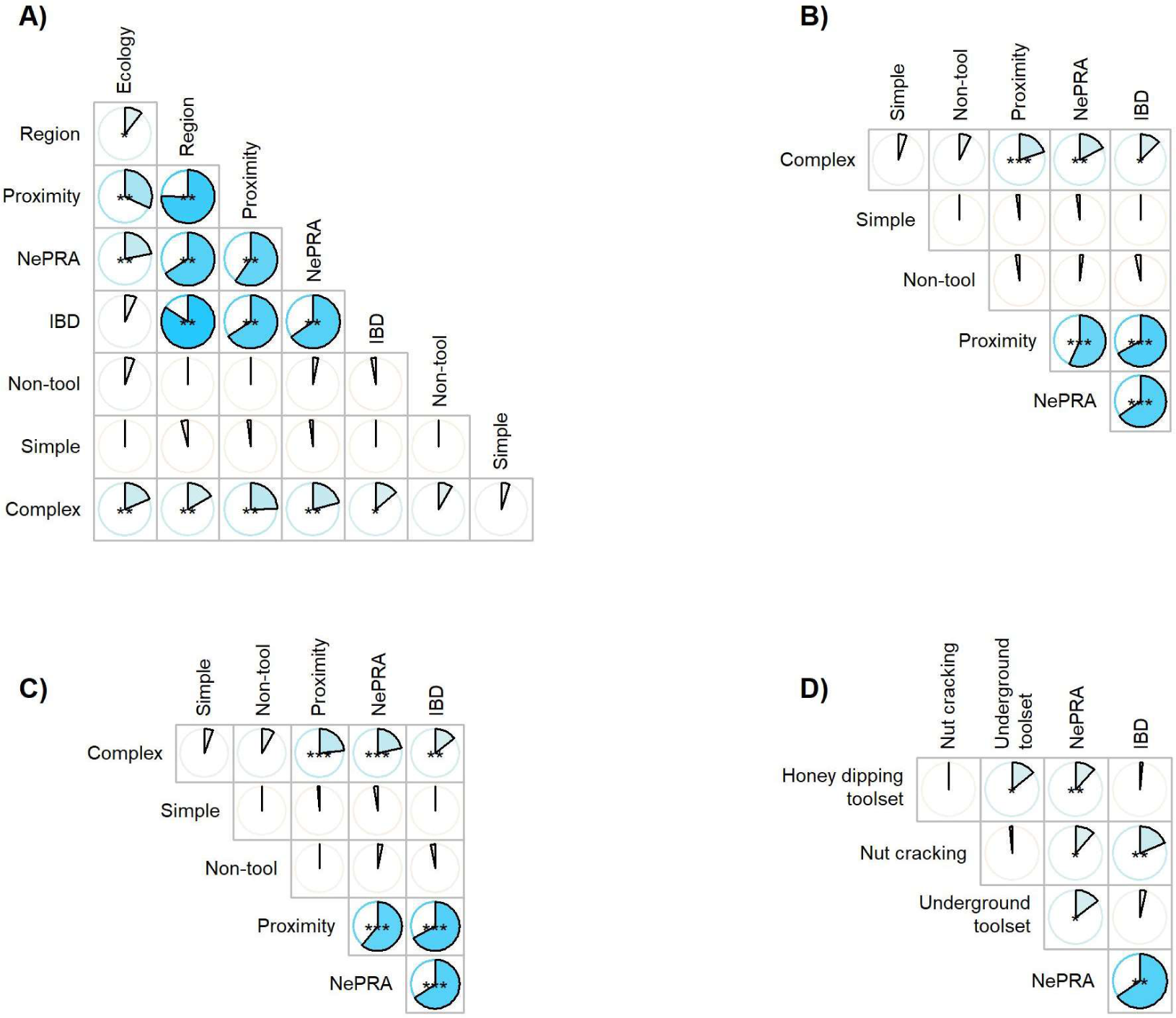
Mantel correlation analysis with 10,000 resamples displaying Spearman correlation coefficients r as the size of the pies and significance showed through the stars (* = 0.05>*P*>0.01, ** 0.01>*P*>0.001, *** P<0.001) testing the null hypothesis that there is no correlation between the respective matrices. Panel A shows correlations from simple Mantel tests which show that while NePRA similarity, IBD sharing, (geographic) proximity and region (subspecies) are highly correlated, all are correlated only with complex tools, apart from NePRA similarity which also has a slight correlation with simple tools. These results correspond to the fits of the Bayesian regressions. Further, partial Mantel tests showed that controlling for habitat (B) or observation time (C) provide similar results. Panel D shows the results from the Mantel tests for each complex trait.

**Fig. S8.**
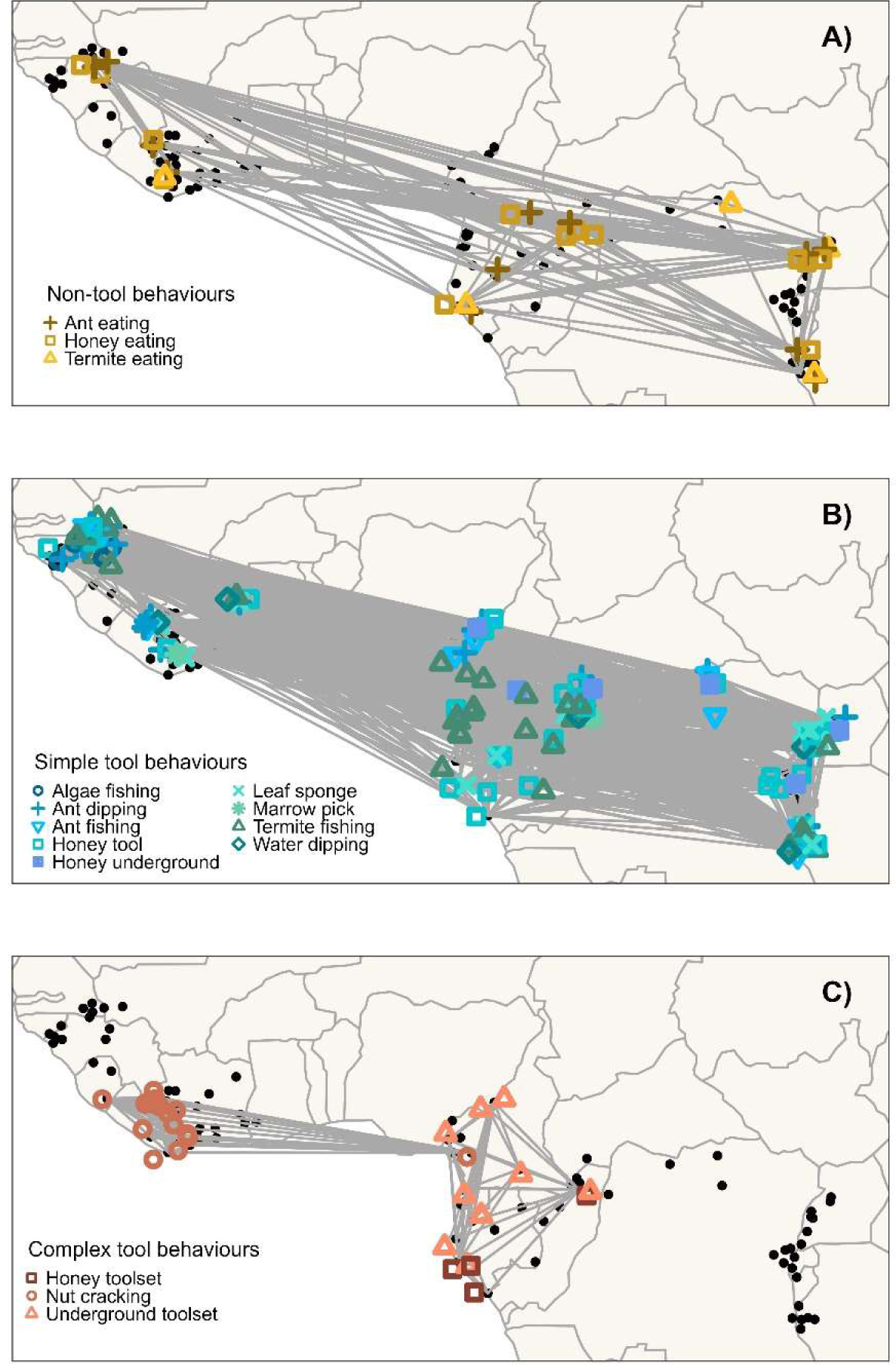
Distribution of the 15 behaviors in all 107 populations from Kalan et al. 2020. Communities (N = 144) at the same site were grouped together into one population.

**Table S1.**
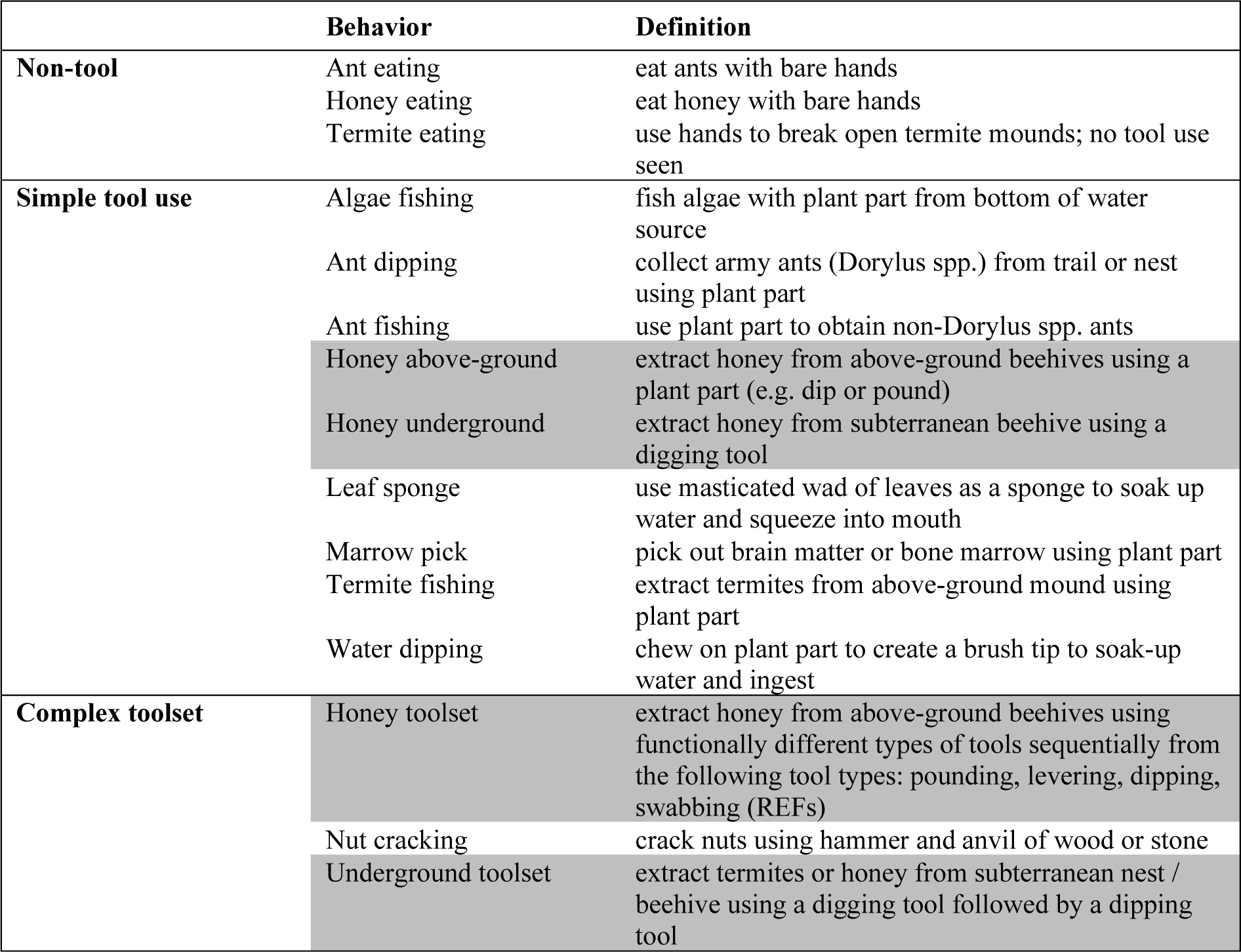
Definitions of the 15 foraging behaviors used in this study and described by Kühl et al. (13). New definitions in this study (highlighted): Honey above-ground, Honey underground, Honey toolset and Underground toolset.

**Table S2.** Results of Bayesian Regression Models for non-tool, simple and complex behaviors. Shared knowledge included as response variable, using weak priors and testing genetic and geographic predictors in separate models. Estimate of the mean marginal posterior distribution, standard deviation of the estimate (sd), the 2.5% and 97.5% credible intervals centered on the mean (CI), proportion of the posterior distribution greater than zero and odds ratios of the mean estimate (OR) are reported. The brackets include the number of dyads. *Average across all six models per response category.

|  | Estimate | sd | CI 2.5% | CI 97.5% | Post. Dist.>0 | OR |
| --- | --- | --- | --- | --- | --- | --- |
| **Non-tool behaviors (n = 595)** |  |  |  |  |  |  |
| Intercept* | -1.832 | 1.223 | -4.378 | 0.433 | 0.059 | 0.160 |
| NePRA binary | 0.009 | 0.401 | -0.767 | 0.800 | 0.514 | 1.009 |
| NePRA similarity | -0.074 | 0.208 | -0.469 | 0.345 | 0.357 | 0.929 |
| IBD binary (n = 561) | -0.388 | 0.452 | -1.309 | 0.491 | 0.197 | 0.678 |
| Distance | 0.241 | 0.233 | -0.202 | 0.727 | 0.856 | 1.273 |
| Same subspecies | 0.044 | 0.419 | -0.801 | 0.833 | 0.547 | 1.045 |
| Same habitat* | 0.539 | 0.406 | -0.258 | 1.344 | 0.909 | 1.717 |
| Shared observation time (months)* | 0.438 | 0.114 | 0.226 | 0.679 | 1.00 | 1.550 |
| **Simple behaviors (n = 595)** |  |  |  |  |  |  |
| Intercept* | -2.780 | 0.792 | -4.119 | -1.014 | 0.002 | 0.062 |
| NePRA binary | 0.529 | 0.163 | 0.211 | 0.848 | 1.00 | 1.697 |
| NePRA similarity | 0.368 | 0.083 | 0.206 | 0.533 | 1.00 | 1.445 |
| IBD binary (n = 561) | 0.321 | 0.18 | -0.043 | 0.675 | 0.96 | 1.379 |
| Distance | -0.103 | 0.085 | -0.267 | 0.065 | 0.113 | 0.902 |
| Same subspecies | 0.022 | 0.178 | -0.326 | 0.372 | 0.549 | 1.022 |
| Same habitat | 0.006 | 0.164 | -0.314 | 0.327 | 0.519 | 1.008 |
| Shared observation time (months)* | 0.30 | 0.062 | 0.181 | 0.424 | 1.00 | 1.351 |
| **Complex behaviors (n = 595)** |  |  |  |  |  |  |
| Intercept* | -2.615 | 1.135 | -4.908 | -0.434 | 0.012 | 0.078 |
| NePRA binary | 1.739 | 0.537 | 0.762 | 2.842 | 1.00 | 5.692 |
| NePRA similarity | 1.148 | 0.304 | 0.567 | 1.744 | 1.00 | 3.152 |
| IBD binary (n = 561) | 0.943 | 0.401 | 0.137 | 1.735 | 0.989 | 2.568 |
| Distance | -0.91 | 0.209 | -1.344 | -0.521 | 0.00 | 0.403 |
| Same subspecies | 1.324 | 0.401 | 0.562 | 2.113 | 1.00 | 3.758 |
| Same habitat | 1.889 | 0.591 | 0.808 | 3.125 | 1.00 | 6.671 |
| Shared observation time (months)* | 0.347 | 0.113 | 0.118 | 0.562 | 0.996 | 1.417 |

**Table S3.**
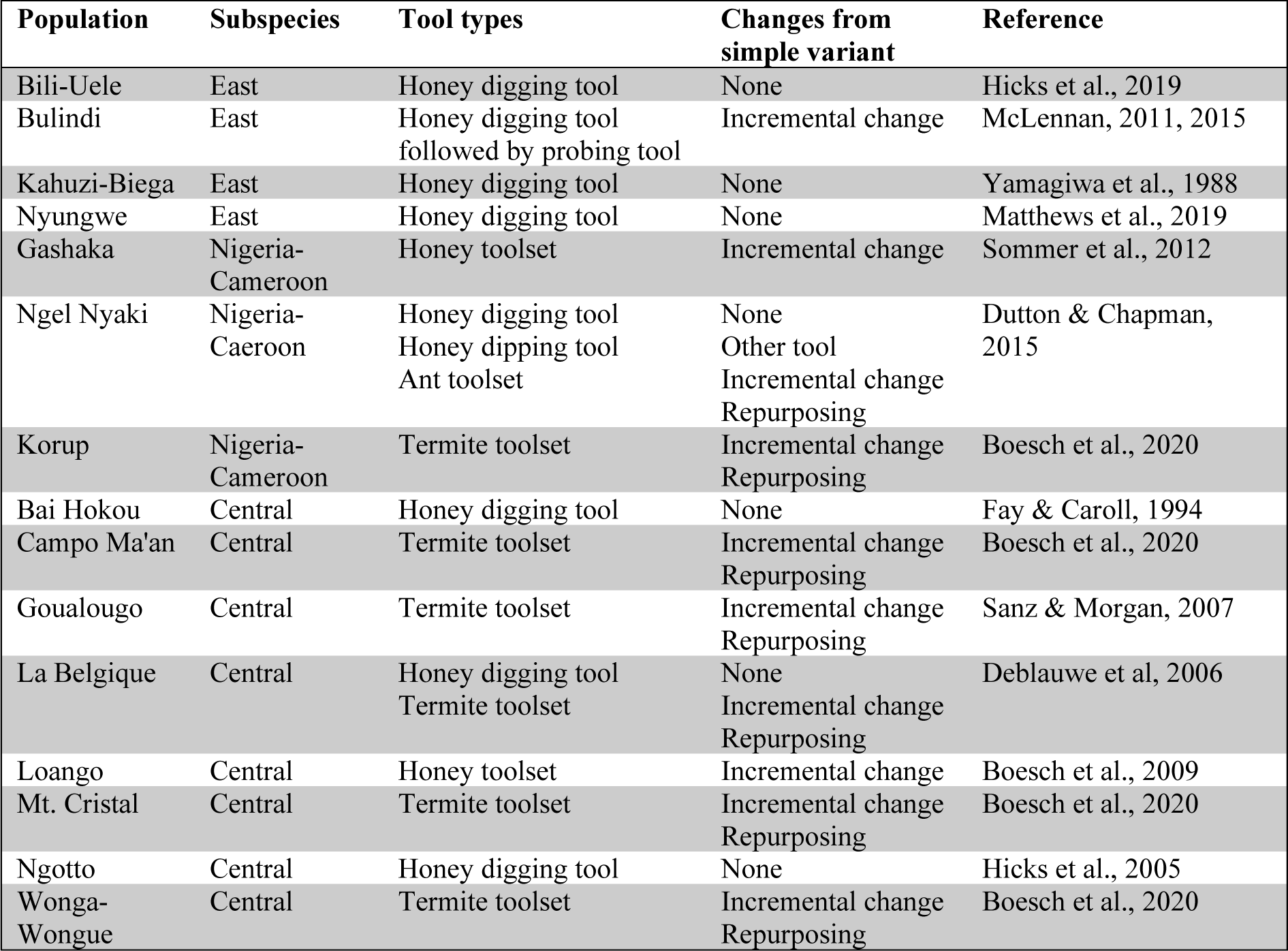
Description of type of underground foraging behaviors at 15 chimpanzee populations. *Honey digging tool*: Underground honey, beehive or bee larvae extraction using a digging tool to enlarge the subterranean beehive entrance; *Honey toolset*: Digging for a beehive followed by the use of a dipping tool to collect honey; *Termite toolset*: Use of a digging tool to perforate the ground and create a tunnel to the subterranean termite mound followed by a fishing tool to collect termites; *Ant toolset*: Digging tool and dipping tool found at the same subterranean ant nest. The changes related to cumulative culture from the simple variant (underground honey digging tool) are described: Incremental change = increase in number of tools involved, Repurposing = change in resource type exploited. The reporting literature was cited.

## References and Notes

1. A. Whiten, C. P. van Schaik, The evolution of animal “cultures” and social intelligence. Philosophical Transactions of the Royal Society B: Biological Sciences 362, 603–620 (2007).

2. F. B. M. de Waal, Animal conformists. Science 340, 437–438 (2013).

3. B. R. Jesmer, J. A. Merkle, J. R. Goheen, E. O. Aikens, J. L. Beck, A. B. Courtemanch, M. A. Hurley, D. E. McWhirter, H. M. Miyasaki, K. L. Monteith, M. J. Kauffman, Is ungulate migration culturally transmitted? Evidence of social learning from translocated animals. Science 361, 1023–1025 (2018).

4. C. Schuppli, C. P. van Schaik, Animal cultures: how we’ve only seen the tip of the iceberg. Evolutionary Human Sciences 1, e2 (2019).

5. A. Whiten, The burgeoning reach of animal culture. Science 372, eabe6514 (2021).

6. J. Henrich, The secret of our success: How culture is driving human evolution, domesticating our species, and making us smarter (Princeton University Press, 2017).

7. A. Mesoudi, A. Thornton, What is cumulative cultural evolution? Proceedings of the Royal Society B: Biological Sciences 285, 20180712 (2018).

8. M. Tomasello, A. C. Kruger, H. H. Ratner, Cultural learning. Behavioral and Brain Sciences 16, 495–552 (1993).

9. C. Tennie, J. Call, M. Tomasello, Ratcheting up the ratchet: on the evolution of cumulative culture. Philosophical Transactions of the Royal Society B: Biological Sciences 364, 2405–2415 (2009).

10. C. Tennie, L. M. Hopper, C. P. van Schaik, “On the origin of cumulative culture: consideration of the role of copying in culture-dependent traits and a reappraisal of the zone of latent solutions hypothesis” in Chimpanzees in context: A comparative perspective on chimpanzee behavior, cognition, conservation, and welfare, L. M. Hopper, S. R. Ross, Eds. (The University of Chicago Press, 2020), pp. 428–453.

11. B. S. Rawlings, C. H. Legare, S. F. Brosnan, G. L. Vale, Leveling the playing field in studying cumulative cultural evolution: conceptual and methodological advances in nonhuman animal research. Journal of Experimental Psychology: Animal Learning and Cognition 47, 252–273 (2021).

12. A. K. Kalan, L. Kulik, M. Arandjelovic, C. Boesch, F. Haas, P. Dieguez, C. D. Barratt, E. E. Abwe, A. Agbor, S. Angedakin, F. Aubert, E. A. Ayimisin, E. Bailey, M. Bessone, G. Brazzola, V. E. Buh, R. Chancellor, H. Cohen, C. Coupland, B. Curran, E. Danquah, T. Deschner, D. Dowd, M. Eno-Nku, J. M. Fay, A. Goedmakers, A.-C. Granjon, J. Head, D. Hedwig, V. Hermans, K. J. Jeffery, S. Jones, J. Junker, P. Kadam, M. Kambi, I. Kienast, D. Kujirakwinja, K. E. Langergraber, J. Lapuente, B. Larson, K. C. Lee, V. Leinert, M. Llana, S. Marrocoli, A. C. Meier, B. Morgan, D. Morgan, E. Neil, S. Nicholl, E. Normand, L. J. Ormsby, L. Pacheco, A. Piel, J. Preece, M. M. Robbins, A. Rundus, C. Sanz, V. Sommer, F. Stewart, N. Tagg, C. Tennie, V. Vergnes, A. Welsh, E. G. Wessling, J. Willie, R. M. Wittig, Y. G. Yuh, K. Zuberbühler, H. S. Kühl, Environmental variability supports chimpanzee behavioural diversity. Nature communications 11, 4451 (2020).

13. H. S. Kühl, C. Boesch, L. Kulik, F. Haas, M. Arandjelovic, P. Dieguez, G. Bocksberger, M. B. McElreath, A. Agbor, S. Angedakin, E. A. Ayimisin, E. Bailey, D. Barubiyo, M. Bessone, G. Brazzola, R. Chancellor, H. Cohen, C. Coupland, E. Danquah, T. Deschner, D. Dowd, A. Dunn, V. E. Egbe, H. Eshuis, A. Goedmakers, A.-C. Granjon, J. Head, D. Hedwig, V. Hermans, I. Imong, K. J. Jeffery, S. Jones, J. Junker, P. Kadam, M. Kambere, M. Kambi, I. Kienast, D. Kujirakwinja, K. E. Langergraber, J. Lapuente, B. Larson, K. Lee, V. Leinert, M. Llana, G. Maretti, S. Marrocoli, R. Martin, T. J. Mbi, A. C. Meier, B. Morgan, D. Morgan, F. Mulindahabi, M. Murai, E. Neil, P. Niyigaba, L. J. Ormsby, R. Orume, L. Pacheco, A. Piel, J. Preece, S. Regnaut, A. Rundus, C. Sanz, J. van Schijndel, V. Sommer, F. Stewart, N. Tagg, E. Vendras, V. Vergnes, A. Welsh, E. G. Wessling, J. Willie, R. M. Wittig, Y. G. Yuh, K. Yurkiw, K. Zuberbühler, A. K. Kalan, Human impact erodes chimpanzee behavioral diversity. Science 363, 1453–1455 (2019).

14. T. Nishida, K. Zamma, T. Matsusaka, A. Inaba, W. C. McGrew, Chimpanzee behavior in the wild: An audio-visual encyclopedia (Springer Science & Business Media, 2010).

15. A. Whiten, J. Goodall, W. C. McGrew, T. Nishida, V. Reynolds, Y. Sugiyama, C. E. G. Tutin, R. W. Wrangham, C. Boesch, Cultures in chimpanzees. Nature 399, 682–685 (1999).

16. A. Whiten, J. Goodall, W. C. McGrew, T. Nishida, V. Reynolds, Y. Sugiyama, C. E. G. Tutin, R. W. Wrangham, C. Boesch, Charting cultural variation in chimpanzees. Behaviour 138, 1481–1516 (2001).

17. C. Boesch, Wild cultures: A comparison between chimpanzee and human cultures (Cambridge University Press, 2012).

18. A. C. Boesch, A. K. Kalan, R. Mundry, M. Arandjelovic, S. Pika, P. Dieguez, E. A. Ayimisin, Barciela, C. Coupland, V. E. Egbe, M. Eno-Nku, J. M. Fay, D. Fine, R. A. Hernandez Aguilar, V. Hermans, P. Kadam, M. Kambi, M. Llana, G. Maretti, D. Morgan, M. Murai, E. Neil, S. Nicholl, L. J. Ormsby, R. Orume, L. Pacheco, A. Piel, C. Sanz, L. Sciaky, F. A. Stewart, N. Tagg, E. G. Wessling, J. Willie, H. S. Kühl, Chimpanzee ethnography reveals unexpected cultural diversity. Nature Human Behaviour 4, 910–916 (2020).

19. W. C. McGrew, Chimpanzee material culture: Implications for human evolution (Cambridge University Press, 1992).

20. W. C. McGrew, Chimpanzee Technology. Science 328, 579–580 (2010).

21. A. B. Migliano, L. Vinicius, The origins of human cumulative culture: from the foraging niche to collective intelligence. Philosophical Transactions of the Royal Society B: Biological Sciences 377, 20200317 (2022).

22. J. Henrich, Demography and cultural evolution: how adaptive cultural processes can produce maladaptive losses: the Tasmanian case. American Antiquity 69, 197–214 (2004).

23. M. Derex, R. Boyd, Partial connectivity increases cultural accumulation within groups. Proceedings of the National Academy of Sciences 113, 2982–2987 (2016).

24. A. B. Migliano, F. Battiston, S. Viguier, A. E. Page, M. Dyble, R. Schlaepfer, D. Smith, L. Astete, M. Ngales, J. Gomez-Gardenes, V. Latora, L. Vinicius, Hunter-gatherer multilevel sociality accelerates cumulative cultural evolution. Science advances 4, eaax5913 (2020).

25. C. Padilla-Iglesias, L. M. Atmore, J. Olivero, K. Lupo, A. Manica, E. A. Isaza, L. Vinicius, A. B. Migliano, Population interconnectivity over the past 120,000 years explains distribution and diversity of Central African hunter-gatherers. Proceedings of the National Academy of Sciences 119, e2113936119 (2022).

26. C. Padilla-Iglesias, J. Blanco-Portillo, A. Ioannidis, A. Manica, L. Vinicius, A. B. Migliano, Cultural evolution of Central African hunter-gatherers reflects a deep history of interconnectivity. [Preprint Version 1] (2022), 10.21203/rs.3.rs-2205369/v1.

27. C. Fontsere, M. Kuhlwilm, C. Morcillo-Suarez, M. Alvarez-Estape, J. D. Lester, P. Gratton, J. M. Schmidt, P. Dieguez, T. Aebischer, P. Álvarez-Varona, A. Agbor, S. Angedakin, A. K. Assumang, E. A. Ayimisin, E. Bailey, D. Barubiyo, M. Bessone, A. Carretero-Alonso, R. Chancellor, H. Cohen, E. Danquah, T. Deschner, A. Dunn, J. Dupain, V. E. Egbe, O. Feliu, A. Goedmakers, A.-C. Granjon, J. Head, D. Hedwig, V. Hermans, R. A. Hernandez-Aguilar, I. Imong, S. Jones, J. Junker, P. Kadam, M. Kaiser, M. Kambere, M. V. Kambale, A. K. Kalan, I. Kienast, D. Kujirakwinja, K. Langergraber, J. Lapuente, B. Larson, A. Laudisoit, K. Lee, M. Llana, M. Llorente, S. Marrocoli, D. Morgan, F. Mulindahabi, M. Murai, E. Neil, S. Nicholl, S. Nixon, E. Normand, C. Orbell, L. J. Ormsby, L. Pacheco, A. Piel, L. Riera, M. M. Robbins, A. Rundus, C. Sanz, L. Sciaky, V. Sommer, F. A. Stewart, N. Tagg, L. R. Tédonzong, E. Ton, J. van Schijndel, V. Vergnes, E. G. Wessling, J. Willie, R. M. Wittig, Y. G. Yuh, K. Yurkiw, K. Zuberbuehler, J. Hecht, L. Vigilant, C. Boesch, A. M. Andrés, D. A. Hughes, H. S. Kühl, E. Lizano, M. Arandjelovic, T. Marques-Bonet, Population dynamics and genetic connectivity in recent chimpanzee history. Cell Genomics 2, 100133 (2022).

28. M. de Manuel, M. Kuhlwilm, P. Frandsen, V. C. Sousa, T. Desai, J. Prado-Martinez, J. Hernandez-Rodriguez, I. Dupanloup, O. Lao, P. Hallast, J. M. Schmidt, J. M. Heredia Genestar, A. Benazzo, G. Barbujani, B. M. Peter, L. F. K. Kuderna, F. Casals, S. Angedakin, M. Arandjelovic, C. Boesch, H. Kühl, L. Vigilant, K. Langergraber, J. Novembre, M. Gut, I. Gut, A. Navarro, F. Carlsen, A. M. Andrés, H. R. Siegismund, A. Scally, L. Excoffier, C. Tyler-Smith, S. Castellano, Y. Xue, C. Hvilsom, T. Marques-Bonet, Chimpanzee genomic diversity reveals ancient admixture with bonobos. Science 354, 477–481 (2016).

29. A. E. Pusey, C. Packer, “Dispersal and philopatry” in Primate societies, B. B. Smuts, D. L. Cheney, R. M. Seyfarth, R. W. Wrangham, Eds. (University of Chicago Press, Chicago, 1986), pp. 250–266.

30. L. V. Luncz, C. Boesch, Tradition over trend: neighboring chimpanzee communities maintain differences in cultural behavior despite frequent immigration of adult females. American Journal of Primatology 76, 649–657 (2014).

31. K. R. Hill, B. M. Wood, J. Baggio, A. M. Hurtado, R. T. Boyd, Hunter-gatherer inter-band interaction rates: implications for cumulative culture. PLoS One 9, e102806 (2014).

32. J. D. Lester, L. Vigilant, P. Gratton, M. S. McCarthy, C. D. Barratt, P. Dieguez, A. Agbor, P. Álvarez-Varona, S. Angedakin, E. A. Ayimisin, E. Bailey, M. Bessone, G. Brazzola, R. Chancellor, H. Cohen, E. Danquah, T. Deschner, V. E. Egbe, M. Eno-Nku, A. Goedmakers, A.-C. Granjon, J. Head, D. Hedwig, R. A. Hernandez-Aguilar, K. J. Jeffery, S. Jones, J. Junker, P. Kadam, M. Kaiser, A. K. Kalan, L. Kehoe, I. Kienast, K. E. Langergraber, J. Lapuente, A. Laudisoit, K. Lee, S. Marrocoli, V. Mihindou, D. Morgan, G. Muhanguzi, E. Neil, S. Nicholl, C. Orbell, L. J. Ormsby, L. Pacheco, A. Piel, M. M. Robbins, A. Rundus, C. Sanz, L. Sciaky, A. M. Siaka, V. Städele, F. Stewart, N. Tagg, E. Ton, J. van Schijndel, M. K. Vyalengerera, E. G. Wessling, J. Willie, R. M. Wittig, Y. G. Yuh, K. Yurkiw, K. Zuberbuehler, C. Boesch, H. S. Kühl, M. Arandjelovic, Recent genetic connectivity and clinal variation in chimpanzees. Communications Biology 4, 283 (2021).

33. S. Carvalho, T. Matsuzawa, W. C. McGrew, “From pounding to knapping: How chimpanzees can help us to model hominin lithics” in Tool use in animals cognition and ecology, C. M. Sanz, J. Call, C. Boesch, eds. (Cambridge University Press, 2010), pp. 225–241.

34. C. Sanz, J. Call, D. Morgan, Design complexity in termite-fishing tools of chimpanzees (Pan troglodytes). Biology Letters 5, 293–296 (2009).

35. G. R. Pradhan, C. Tennie, C. P. van Schaik, Social organization and the evolution of cumulative technology in apes and hominins. Journal of human evolution 63, 180–190 (2012).

36. F. Battiston, V. Nicosia, V. Latora, Structural measures for multiplex networks. Physical Review E 89, 032804 (2014).

37. S. J. Lycett, M. Collard, W. C. McGrew, Are behavioral differences among wild chimpanzee communities genetic or cultural? An assessment using tool-use data and phylogenetic methods. American Journal of Physical Anthropology 142, 461–467 (2010).

38. R. Kendal, L. M. Hopper, A. Whiten, S. F. Brosnan, S. P. Lambeth, S. J. Schapiro, W. Hoppitt, Chimpanzees copy dominant and knowledgeable individuals: implications for cultural diversity. Evolution and Human Behavior 36, 65–72 (2015).

39. M. S. Granovetter, The strength of weak ties. American Journal of Sociology 78, 1360–1380 (1973).

40. C. Hobaiter, T. Poisot, K. Zuberbühler, W. Hoppitt, T. Gruber T. Social network analysis shows direct evidence for social learning of tool use in wild chimpanzees. PLOS Biology 12, e1001960 (2014).

41. C. P. van Schaik, G. R. Pradhan, C. Tennie, Teaching and curiosity: sequential drivers of cumulative cultural evolution in the hominin lineage. Behavioural ecology and sociobiology 73 (2019), pp. 1–11.

42. A. Whiten, “Social learning and culture in the great apes” in The Oxford Handbook of Cultural Evolution, J. Tehrani, R. Kendal, J. Kendal, Eds. (Oxford University Press, 2023), pp. C23S1–C23S31.

43. J. Mercader, H. Barton, J. Gillespie, J. Harris, S. Kuhn, R. Tyler, C. Boesch, 4,300-Year-old chimpanzee sites and the origins of percussive stone technology. Proceedings of the National Academy of Sciences 104, 3043–3048 (2007).

44. A. B. Migliano, A. E. Page, J. Gómez-Gardeñes, G. D. Salali, S. Viguier, M. Dyble, J. Thompson, N. Chaudhary, D. Smith, J. Strods, R. Mace, M. G. Thomas, V. Latora, L. Vinicius, Characterization of hunter-gatherer networks and implications for cumulative culture. Nature Human Behaviour 1, 0043 (2017).

45. M. Dyble, G. D. Salali, N. Chaudhary, A. Page, D. Smith, J. Thompson, L. Vinicius, R. Mace, A. B. Migliano, Sex equality can explain the unique social structure of hunter-gatherer bands. Science 348, 796–798 (2015).

46. M. Dyble, J. Thompson, D. Smith, G. D. Salali, N. Chaudhary, A. E. Page, L. Vinicius, R. Mace, A. B. Migliano, Networks of food sharing reveal the functional significance of multilevel sociality in two hunter-gatherer groups. Current Biology 26, 2017–2021 (2016).

47. A. Powell, S. Shennan, M. G. Thomas, Late Pleistocene demography and the appearance of modern human behavior. Science 324, 1298–1301 (2009).

48. M. Derex, C. Perreault, R. Boyd, Divide and conquer: intermediate levels of population fragmentation maximize cultural accumulation. Philosophical Transactions of the Royal Society B: Biological Sciences 373, 20170062 (2018).

49. P. M. Kappeler, C. P. van Schaik, Evolution of Primate Social Systems. International journal of primatology 23, 707–740 (2002).

50. J. M. Burkart, S. B. Hrdy, C. P. van Schaik, Cooperative breeding and human cognitive evolution. Evolutionary Anthropology 18, 175–186 (2009).

51. A. E. Page, N. Chaudhary, S. Viguier, M. Dyble, J. Thompson, D. Smith, G. D. Salali, R. Mace, A. B. Migliano, Hunter-gatherer social networks and reproductive success. Scientific Reports 7, 1153 (2017).

52. N. Blegen, The earliest long-distance obsidian transport: Evidence from the ∼200 ka Middle Stone Age Sibilo School Road Site, Baringo, Kenya. Journal of Human Evolution 103, 1–19 (2017).

53. A. S. Brooks, J. E. Yellen, R. Potts, A. K. Behrensmeyer, A. L. Deino, D. E. Leslie, S. H. Ambrose, J. R. Ferguson, F. D’Errico, A. M. Zipkin, S. Whittaker, J. Post, E. G. Veatch, K. Foecke, J. B. Clark, Long-distance stone transport and pigment use in the earliest Middle Stone Age. Science 360, 90–94 (2018).

54. E. M. L. Scerri, L. Chikhi, M. G. Thomas, Beyond multiregional and simple out-of-Africa models of human evolution. Nature ecology & evolution 3, 1370–1372 (2019).

55. K. N. Laland, Gene-Culture Coevolution. Encyclopedia of Cognitive Science. 2, 268–274 (2003).

56. C. M. Sanz, D. B. Morgan, Chimpanzee tool technology in the Goualougo Triangle, Republic of Congo. Journal of Human Evolution 52, 420–433 (2007).

57. B. L. Browning, S. R. Browning, Detecting identity by descent and estimating genotype error rates in sequence data. The American Journal of Human Genetics 93, 840–851 (2013).

58. E. A. Thompson, Identity by descent: Variation in meiosis, across genomes, and in populations. Genetics 194, 301–326 (2013).

59. S. Schiffels, W. Haak, P. Paajanen, B. Llamas, E. Popescu, L. Loe, R. Clarke, A. Lyons, R. Mortimer, D. Sayer, C. Tyler-Smith, A. Cooper, R. Durbin, Iron Age and Anglo-Saxon genomes from East England reveal British migration history. Nature communications 7, 10408 (2016).

60. R Core Team, A language and environment for statistical computing. R Foundation for Statistical Computing, Vienna, Austria (2022; https://www.R-project.org/).

61. Posit team, RStudio: Integrated Development Environment for R. Posit Software, PBC, Boston, MA (2023; http://www.posit.co/).

62. H. Schielzeth, Simple means to improve the interpretability of regression coefficients. Methods in Ecology and Evolution 1, 103–113 (2010).

63. P.-C. Bürkner, brms: An R package for Bayesian multilevel models using Stan. Journal of statistical software 80, 1–28 (2017).

64. A. Gelman, J. B. Carlin, H. S. Stern, Bayesian data analysis 3rd edition (CRC Press, 2014).

65. J. Oksanen, G. L. Simpson, F. G. Blanchet, R. Kindt, P. Legendre, P. R. Minchin, R. B. O’Hara, P. Solymos, M. H. H. Stevens, E. Szoecs, H. Wagner, M. Barbour, M. Bedward, B. Bolker, D. Borcard, G. Carvalho, M. Chirico, M. de Caceres, S. Durand, H. B. Antoniazi Evangelista, R. FitzJohn, M. Friendly, B. Furneaux, G. Hannigan, M. O. Hill, L. Lahti, D. McGlinn, M.-H. Ouellette, E. Ribeiro Cunha, T. Smith, A. Stier, C. J.F. ter Braak, Vegan: community ecology package, version 2.4-3. CRAN (2022; https://CRAN.R-project.org/package=vegan).

66. S. C. Goslee, D. L. Urban, The ecodist package for dissimilarity-based analysis of ecological data. Journal of Statistical Software (2007; http://www.jstatsoft.org/).

